# Deciphering the role of *Leptospira* surface protein LigA in modulating the host innate immune response

**DOI:** 10.1101/2020.10.12.329177

**Authors:** Ajay Kumar, Vivek P. Varma, Kavela Sridhar, Mohd Abdullah, Pallavi Vyas, Ashiq T Muhammed, Yung-Fu Chang, Syed M. Faisal

## Abstract

*Leptospira*, a zoonotic pathogen, is known to infect various hosts and can establish persistent infection. This remarkable ability of bacteria is attributed to its potential to modulate the host immune response by exploiting its surface proteins. We have identified and characterized the domain of the variable region of *Leptospira* immunoglobulin-like protein A (LAV) involved in immune modulation. The 11^th^ domain (A_11_) of the variable region of LigA (LAV) induces a strong TLR4 dependent innate response leading to subsequent induction of humoral and cellular immune responses in mice. A_11_ is also involved in acquiring complement regulator FH and binds to host protease Plasminogen (PLG), there by mediating functional activity to escape from complement-mediated killing. The deletion of A_11_ domain significantly impaired TLR4 signaling and subsequent reduction in the innate and adaptive immune response. It also inhibited the binding of FH and PLG thereby mediating killing of bacteria. Our study discovered an unprecedented role of LAV as a nuclease capable of degrading Neutrophil Extracellular Traps (NETs). This nuclease activity was primarily mediated by A_11_. These results highlighted the moonlighting function of LigA and demonstrated that a single domain of a surface protein is involved in evading a myriad of host innate immune defenses, which might allow the persistence of *Leptospira* in different hosts for a long term without clearance.

## Introduction

Leptospirosis is one of the most widespread bacterial zoonosis, particularly in developing countries like India, and one of the major neglected infectious diseases globally^1^. It is caused by the pathogenic spirochete of the genus *Leptospira* that can cause fatal infections involving multiple organs in human and animal hosts. According to WHO, there is a substantial economic burden of human leptospirosis with an estimated 1.03 million cases and 58,900 deaths worldwide annually^2^. The actual burden may be much higher as a lot of cases are not reported due to difficulties associated with diagnosis^2^. The major challenge in combating this zoonosis has been the unavailability of early diagnostics and potent vaccines that can induce cross-protection against various serovars ^3^. Understanding how *Leptospira* escapes from host innate immune defenses to disseminate and colonize in multiple organs for establishing infection will aid in devising prophylactic strategies.

Innate immune responses comprising of soluble factors like antimicrobial peptides and complement proteins, pattern recognition receptors like Toll-like receptors (TLRs) and NOD-like receptors (NLRs), and phagocytic cells such as Dendritic cells (DCs), neutrophils, and macrophages contribute to the killing and removal of invading pathogens by a variety of mechanisms^4^. Signaling through TLRs induces activation of innate immune cells leading to secretion of pro-inflammatory cytokines (IL-6, TNF-α) and expression of surface molecules (CD80, CD86, MHC-II), thereby enabling these cells to become efficient in subsequent activation of adaptive response^5,6^. TLRs play a key role in promoting adaptive immune responses and are also essential for T-cell expansion, differentiation, and memory formation^7^. The Complement system is a vital part of innate immune defense that promptly kills the invading pathogen by opsonization and target lysis^8^. To prevent damage to the host cells, the complement system is tightly regulated by soluble plasma proteins like Factor H (FH) and C4b-binding protein (C4BP)^9^. FH and C4BP regulate the Alternative pathway (AP), Classical pathway (CP), and Lectin pathway of complement activation. Plasmin, the enzymatically active form of plasminogen (PLG) acts as a protease that potentially cleaves complement factors C3b, C4b and C5^10^. Neutrophils are major phagocytic cells that utilize a combination of reactive oxygen species (ROS), cytotoxic granules, antimicrobial peptides, and Neutrophil Extracellular Traps (NETs) to kill and degrade the invading pathogen^11^. However, pathogens have devised several strategies to escape from host innate immune defenses through a mechanism mediated by their surface proteins^12^. These proteins may be pro-inflammatory where they can activate APCs like macrophages and DCs but might also enable the pathogen to avoid recognition through innate receptors (TLRs) through downregulation of their expression or causing antigenic variations to evade from host defences^13,14^. Pathogens escape from complement-mediated killing by expressing surface proteins that acquire complement regulators like FH and C4BP, act as proteases or acquire host proteases that can cleave complement components^8,15^. They may avoid killing by phagocytes like neutrophils by expressing surface proteins, which may help inevading extravasation and chemotaxis, preventing opsonization and phagocytosis, promoting survival inside the neutrophil, and inducing apoptosis or cell death and degrading NETs by virtue of their nuclease activity^16,17^.

Like other pathogens, *Leptospira* has also evolved strategies to modulate the host’s innate immune response by exploiting the capacities of its surface proteins to favor their pathogenesis^18–20^. Toll-like receptors like TLR2 and TLR4 play a major role in host defense as mice lacking these receptors were highly susceptible to *Leptospira* infection^21^. These bacteria likely modulate the expression of surface molecules (proteins, LPS) to avoid recognition through protective TLR2 and TLR4 and establish infection in the host. Several surface proteins of *Leptospira* have been identified as a potent activator of pro-inflammatory response via signaling through both TLR2 and TLR4^22–24^. Besides that, several proteins have been shown to acquire FH, C4BP and PLG on their surface or act as proteases to cleave complement components to evade killing^25–30^. *Leptospira* is known to induce NET; hence it is likely that it might express surface proteins/nucleases like other bacteria to evade NETosis^31^. Thus, identification and characterization of a surface protein involved in the modulation of the host innate immune response will aid in designing a better strategy to combat this bacterial zoonosis.

*Leptospira* immunoglobulin-like (Lig) proteins (LigA and LigB) are surface proteins having 12-13 immunoglobulin-like repeat domains similar to an invasin of *Yersinia* and intimin of *E.coli^32^*. The N terminal region of LigA and LigB from domains 1 to 6.5 are conserved, whereas C terminal regions from domains 6.5 to 13 are variable^32,33^. Lig proteins are expressed during infection and have been shown to bind to multiple components of the host extracellular matrix (ECM), thereby mediating attachment to host cells ^34,35^. They are the most promising vaccine candidate identified to date. Moreover, the variable region of LigA (LAV) comprising domain 6.5-13 (LAV) was shown to be sufficient to induce protection against challenge in the hamster model ^36–41^. Despite various reports confirming the protective role of LAV, its involvement in the modulation of the host innate immune response has not been studied extensively. Several groups demonstrated that Lig proteins bind to FH and C4BP to inhibit lectin, classical and alternative pathways; however, specific domains involved in binding to these regulators have not been characterized^27,29,30,42,43^. Further, the role of the protein in activation of the innate response or evasion from killing by phagocytes has not been reported so far. In the present study, we have demonstrated the role of LAV in modulating the host innate immune response. Using various assays, we identified the domain/s involved in activating of innate and subsequent adaptive immune response and evasion from complement-mediated killing via binding to FH and host protease PLG. Further, we demonstrated LAV’s nuclease activity which might play a major role in evasion from Neutrophil extracellular traps (NETosis).

## Results

### LAV induced TLR4 dependent activation of mouse macrophages

To test whether the immunogenicity of Variable region of A (LAV) correlates to innate immune response activation, we tested its ability to activate mouse macrophages. We cloned, expressed, and purified LAV in a soluble form (Sup Fig. 1). We stimulated mouse macrophages with varying doses (1, 2, and 5μg/ml) of the protein and our result shows that LAV induced production of pro-inflammatory cytokines (IL-6, TNF-α) in dose-dependent manner (Fig. 1A). Taking into account that the purified protein might have LPS contamination, they were pre-treated twice with Polymyxin-B Agarose to remove the endotoxin activity. 500ng/ml LPS pre-incubated with the same concentrations of PMB-agarose was used as control. The concentration of LPS in final protein preparation varied from (0.10–0.15ng/ml). The effect was protein-specific because Proteinase-K plus heating abolished cytokine production (Sup Fig. 2). Besides, PMB inhibited the LPS induced cytokine production but did not attenuate the levels induced by LAV, indicating that the stimulatory effects observed were specific to protein and not due to contamination with LPS (Sup Fig. 2). Next, we tested whether this LAV-induced activation was via signaling through TLR2 or TLR4. As confirmed by confocal microscopy, LAV showed binding, specifically with TLR4 and failed to bind to the TLR2 receptor (Fig. 1B). To verify that this binding leads to activation and subsequent cytokine production, we stimulated WT, TLR2KO, TLR4KO and DKO macrophages and HEK-293T cells expressing these receptors with LAV. Our result shows that while WT and TLR2KO macrophages cells induced significant levels of IL-6 and TNF-α, TLR4KO and DKO macrophages failed to induce these cytokines (Fig. 1C).

**Fig.1:**
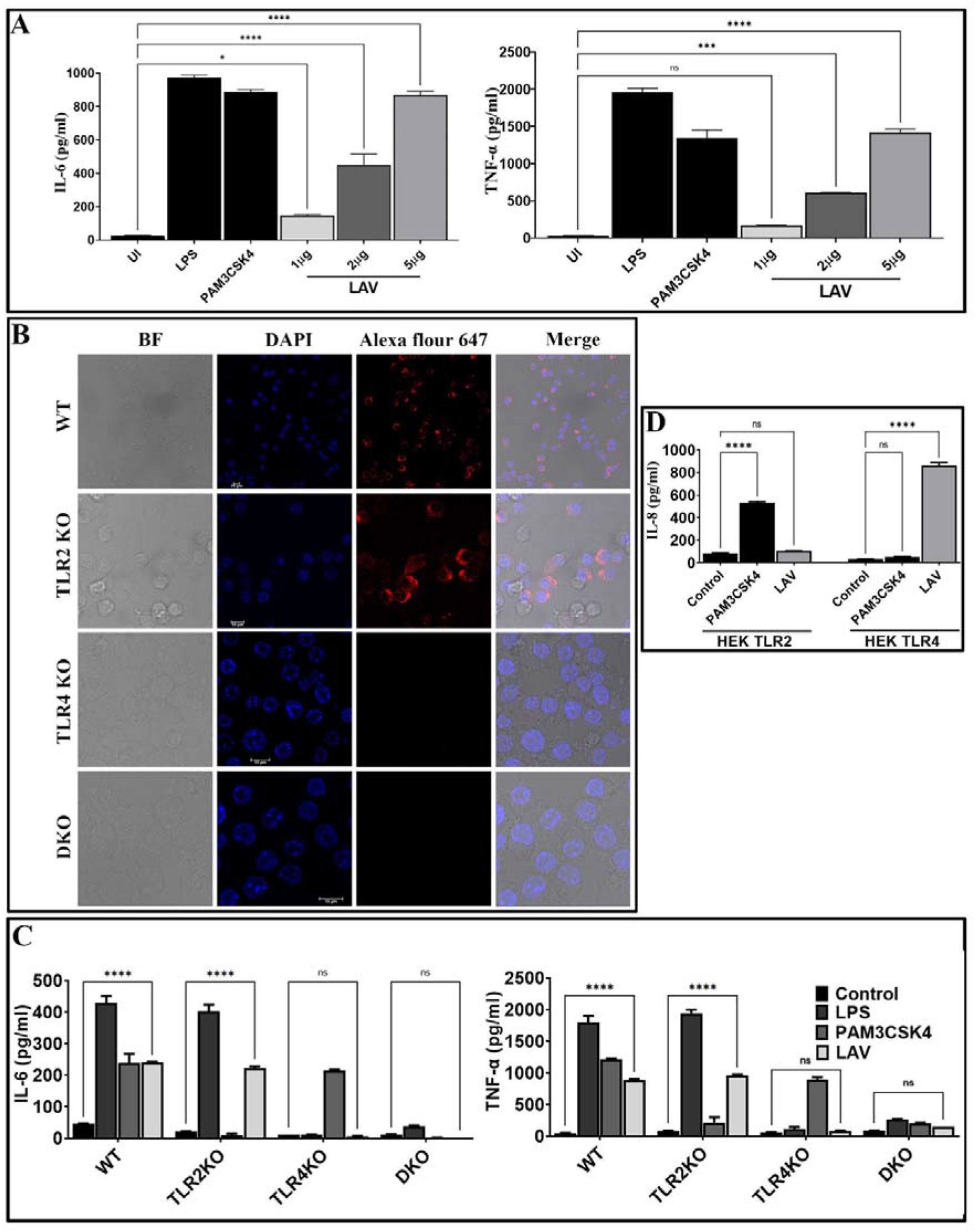
Variable region of LigA (LAV) induced TLR4 dependent activation of mouse macrophages. (**A**) *Screening of pro-inflammatory response of LAV in RAW264.7 cells*. RAW264.7 cell lines were stimulated with LAV in varying concentrations 1 or 2 or 5μg/ml along with PMB for 24h and supernatant was collected to measure levels of TNF-α and IL-6 cytokines by using ELISA. **(B)** *Binding of LAV with TLR4*. WT, TLR2KO, TLR4KO, or DKO macrophages were incubated with LAV (2μg/ml) for 30 min. After washing, cells were fixed and stained with then stained with Alexa Flour 647 conjugated rabbit anti-mouse IgG and analysed by the confocal microscope as described in materials and methods. **(C)** *Pro-inflammatory response of mouse macrophages stimulated with rLig proteins*. WT, TLR2KO, TLR4KO and DKO macrophages cell lines were treated LAV at 2μg/ml for 24h at 37°C in presence of 5%CO_2_ and levels of IL-6 and TNF-α in the supernatants were measured by ELISA. **(D)** *IL-8 response in HEK293-TLR4 cells stimulated with LAV*. HEK293T cells transfected with TLR2, TLR4 and NF-kB reporter plasmids were stimulated with rLAV (2μg/ml) for 24h and IL-8 was measured in the culture supernatant by ELISA. *E. coli* LPS (500ng/ml) or PAM3CSK4(20ng/ml) as TLR4 and TLR2 ligands respectively were used as positive controls in all experiments wherever indicated. Data are representative of three different experiments. Significant differences were calculated using the one or two way ANOVA(****, ***, **, * indicates P < 0.0001, < 0.001, P < 0.01 and P□<□0.05 respectively).

Similarly, HEK-TLR4 stimulated with LAV produced significant levels of IL-8, whereas HEK-TLR2 cells didn’t produce a significant level of this cytokine (Fig. 1D). These results indicate that LAV is a TLR4 ligand that induces signalling through this receptor for the production of pro-inflammatory cytokines.

### 11^th^ domain of the variable region of LigA (A_11_) is involved in signalling through TLR4 for the activation and maturation of macrophages

Since LAV induced TLR4-dependent activation of mouse macrophages, we aimed to identify and characterize the domain/s involved in activation. We cloned, expressed and purified the individual domain (A_8_-A_13_) and tested their ability to activate mouse macrophages (RAW264.7 cells). Our result shows that only 11^th^ domain (A_11_) induced a significant level of IL-6 and TNF-α (Fig.2A). To confirm that A_11_ is involved in the production of cytokines, we created domain deletion mutants of LAV(AΔ_8_-AΔ_13_) and purified the proteins in the soluble form (Sup Fig.1B). We tested the ability of these mutants to activate mouse macrophages, and our result shows that all the deletion mutants of LAV induced production of IL-6 and TNF-α except AΔ_11_, further confirming that this domain is involved in the activation of macrophages and subsequent production of cytokines (Fig. 2B). We tested its binding with the receptor to confirm that A_11_ is involved in interaction and subsequent signaling via TLR4. Confocal microscopy confirmed the binding of A_11_ with the mouse TLR4 as strong anti-A_11_ fluorescence was observed on the surface of WT and TLR2KO cells but little fluorescence on TLR4KO or DKO cells. Further, there was very little anti-AΔ_11_ fluorescence on the surface of all cell types indicating that this protein failed to bind to the TLR receptor (Fig. 2C). To confirm that this TLR4 binding leads to activation of these cells, we stimulated mouse WT, TLR2KO, TLR4KO, and DKO macrophages with LAV, A_11_, and AΔ_11_, and our results indicate that A_11_ induced IL-6 and TNF-α production via signaling through TLR4 as TLR4KO and DKO macrophages failed to induce any significant level of these cytokines. Further, the inability of AΔ_11_ to induce substantial levels of cytokines in WT or TLR2 KO macrophages indicates that the 11^th^ domain is critical for signaling via TLR4 (Fig. 2D). To confirm whether stimulation with A_11_ causes macrophage activation and maturation, we analyzed the expression of costimulatory molecules (CD80, CD86, and CD40) and the maturation marker (MHC-II) in RAW264.7 cells. Our Flow cytometry results show that LAV and A_11_ significantly enhanced the expression of CD80, CD86, CD40, and MHCII, whereas AΔ_11_ failed to upregulate them, indicating that this domain is involved in enhancing the expression of these surface molecules (Fig. 2E). To understand whether deletion of the 11^th^ domain leads to structural changes in the protein, which might be contributing to impairing its innate immune activity, we did CD spectroscopy and our result shows that deletion of A_11_ has reduced the helix and beta sheets and, in turn, distorted the structure but didn’t have any effect on proper folding of LAV (Sup. Fig. 3). In conclusion, our results demonstrate that A_11_ is involved in TLR4 dependent activation and maturation of mouse macrophages.

**Fig.2:**
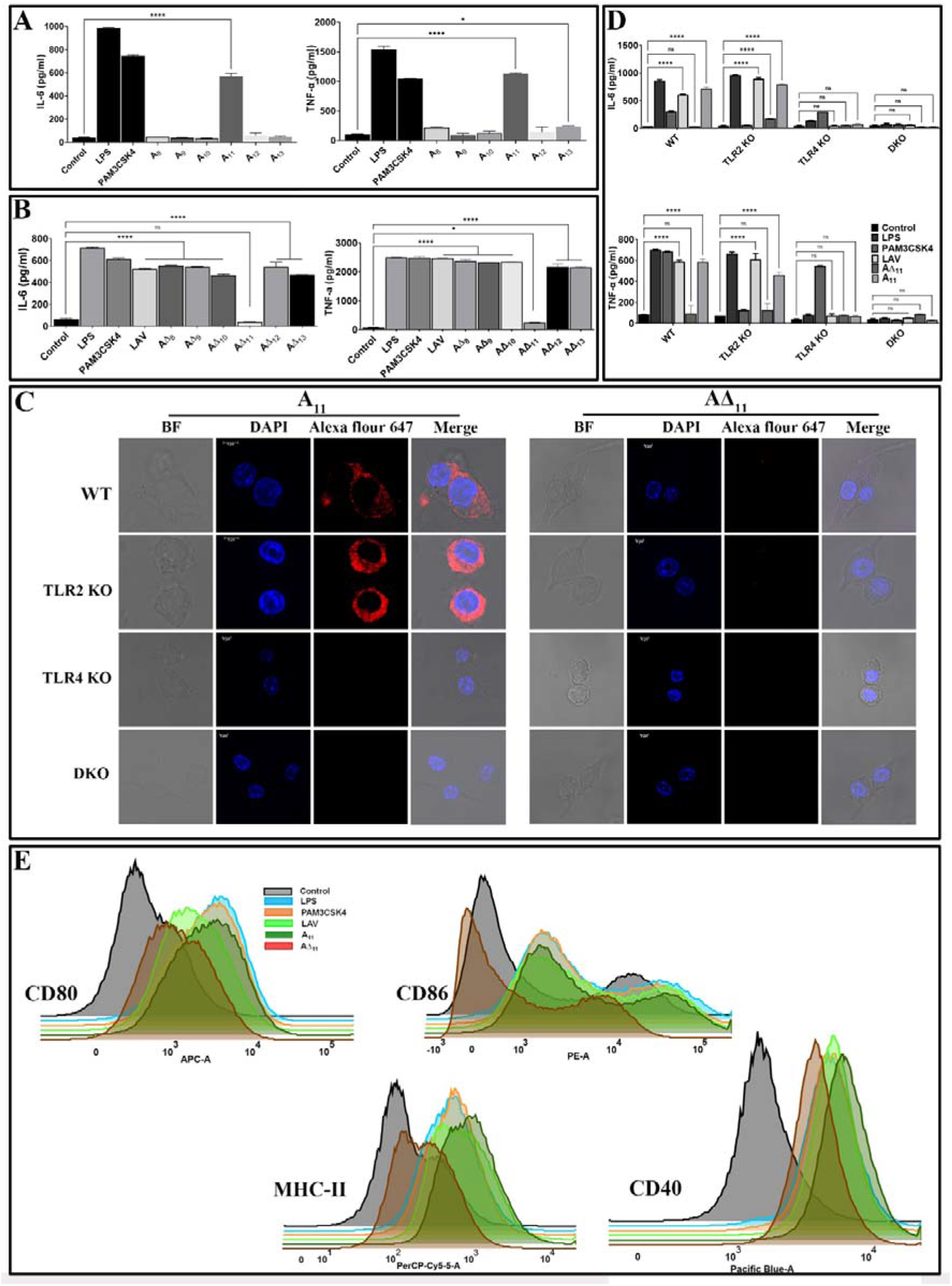
11^th^ domain of LigA (A_11_) is involved in binding to TLR4 and subsequent signalling leading to activation and maturation of mouse macrophages. **(A)** *Screening of pro-inflammatory response of individual domains of LAV in RAW264.7 cells* RAW264.7 cells were incubated with individual domains (A_8_-A_13_) at concentration of 2μg/ ml pre-treated with Polymixin B for 24h at 37°C in presence of 5%CO_2_ and supernatant was collected to measure levels of IL-6 and TNF-α by ELISA. **(B)** *Screening of pro-inflammatory response of domain deletion mutants of LAV in RAW264.7 cells*. RAW264.7 cells were stimulated with LAV or corresponding deletion mutants (AΔ_8_-AΔ_13_) at a concentration of 2μg/ml pre-treated with Polymixin B for 24h at 37°C in the presence of 5%CO_2_ and supernatant was collected to measure levels of IL-6 and TNF-α by ELISA. **(C)***Binding of A_11_ with the mouse TLR4*. WT, TLR2KO, TLR4KO or DKO macrophages cell lines were incubated with A_11_ and AΔ_11_ (2μg/ml) for 30 min. After washing, cells were fixed and stained with respective antibodies and analysed by confocal microscope as described in materials and methods. **(D)** *The pro-inflammatory response of mouse macrophages stimulated with A_11_*. WT, TLR2KO, TLR4KO and DKO bone marrow-derived macrophages cell lines were treated with LAV, A_11_, AΔ_11_ (2μg/ml), LPS (500ng/ml) or PAM3CSK4(20ng/ml) for 24h at 37°C in presence of 5%CO_2_ and levels of IL-6 and TNF-α in the supernatants were measured with ELISA. **(E)** *A_11_ enhanced the expression of surface markers in mouse macrophages. RAW264.7* cells were incubated with LPS (500 ng/ml) or PAM3CSK (20ng/ml) or LAV or A_11_ or AΔ_11_ (2μg/ml) for 24h at 37°C in presence of 5%CO_2_. Cells were stained with fluorochrome-conjugated antibodies and then analyzed by Flow cytometry as described in materials and methods. Control indicates uninduced or unstimulated cells, *E. coli* LPS (500ng/ml) or PAM3CSK4(20ng/ml) as TLR4 and TLR2 ligands respectively were used as positive controls in all experiments wherever indicated. Data are representative of three different experiments. Significant differences were calculated using the one or two way ANOVA(****, ***, **, * indicates P < 0.0001, < 0.001, P < 0.01 and P□<□0.05 respectively).

### A_11_ induces immune activation via MAPK signaling involving the MyD88 adapter

Since TLR4 involves both MyD88 and TRIF adapter for downstream signaling and A_11_ induced immune activation through TLR4, we examined the adapter molecule involved in the signaling. We stimulated MyD88KO, TRIFKO, and TMDKO macrophages with A_11_, and our results show that the signaling pathway involves MyD88 adapter as MyD88KO macrophages failed to induce significant levels of IL-6 and TNF-α. In contrast, there was no difference in the production of these cytokines in TRIFKO macrophages (Fig. 3A). Because MAPKs are critical factors involved in cellular responses to inflammatory stimuli, we examined the activation of this pathway in response to A_11_. We stimulated mouse WT, TLR2KO, TLR4KO, and DKO macrophages with A_11_ and analyzed the phosphorylation of P38, JNK, ERK and degradation of IkBα (Fig. 3B). Next, to elucidate the functional role of these kinases in A_11_ induced macrophage activation and maturation, we used pharmacological inhibitors of these pathways and analyzed cytokines in RAW264.7 cells pre-treated with or without inhibitors of NF-kB, JNK, p38MAPK or ERK. IL-6 and TNF-α production was significantly blocked by p38 inhibitor (P <0.05, 50% inhibition with 2μg/ml A_11_) and by JNK and NF-kB (P, <0.05, 30% inhibition with 2μg/ml A_11_). ERK inhibitor didn’t effect the production of cytokine, indicating that this pathway is not involved in signalling. The production of TNF-α was also significantly blocked by JNK, p38, and NF-κB inhibitor (P <0.05, 60% inhibition). A combination of all inhibitors completely inhibited A_11_ induced cytokine production (Fig. 3C). All these results suggest that A_11_ stimulates the production of pro-inflammatory cytokines through p38, JNK and NF-kB pathways. The ability of A_11_ to regulate innate responses was further investigated based on the expression of key inflammatory cytokine and chemokine genes at various time points (4, 24, and 48h). WT, TLR2KO, TLR4KO and DKO mouse macrophage were stimulated with A_11_, and expression of mRNA transcript was analyzed by RT-PCR. A_11_ induced CXCL10, IL-1β, TNF-α, COX2, iNOS, MCP-1, and IL-6 in WT and TLR2KO mouse macrophage at 4h and 24h time point which was significantly reduced or down-regulated in TLR4KO and DKO macrophages (Fig. 3D). PAM3CSK4 (TLR2 ligand) and LPS (TLR4 ligand) showed significant upregulation of genes involved in TLR2 and TLR4 signalling as expected. These results demonstrate that A_11_ induced TLR4 dependent expression of innate response genes.

**Fig.3:**
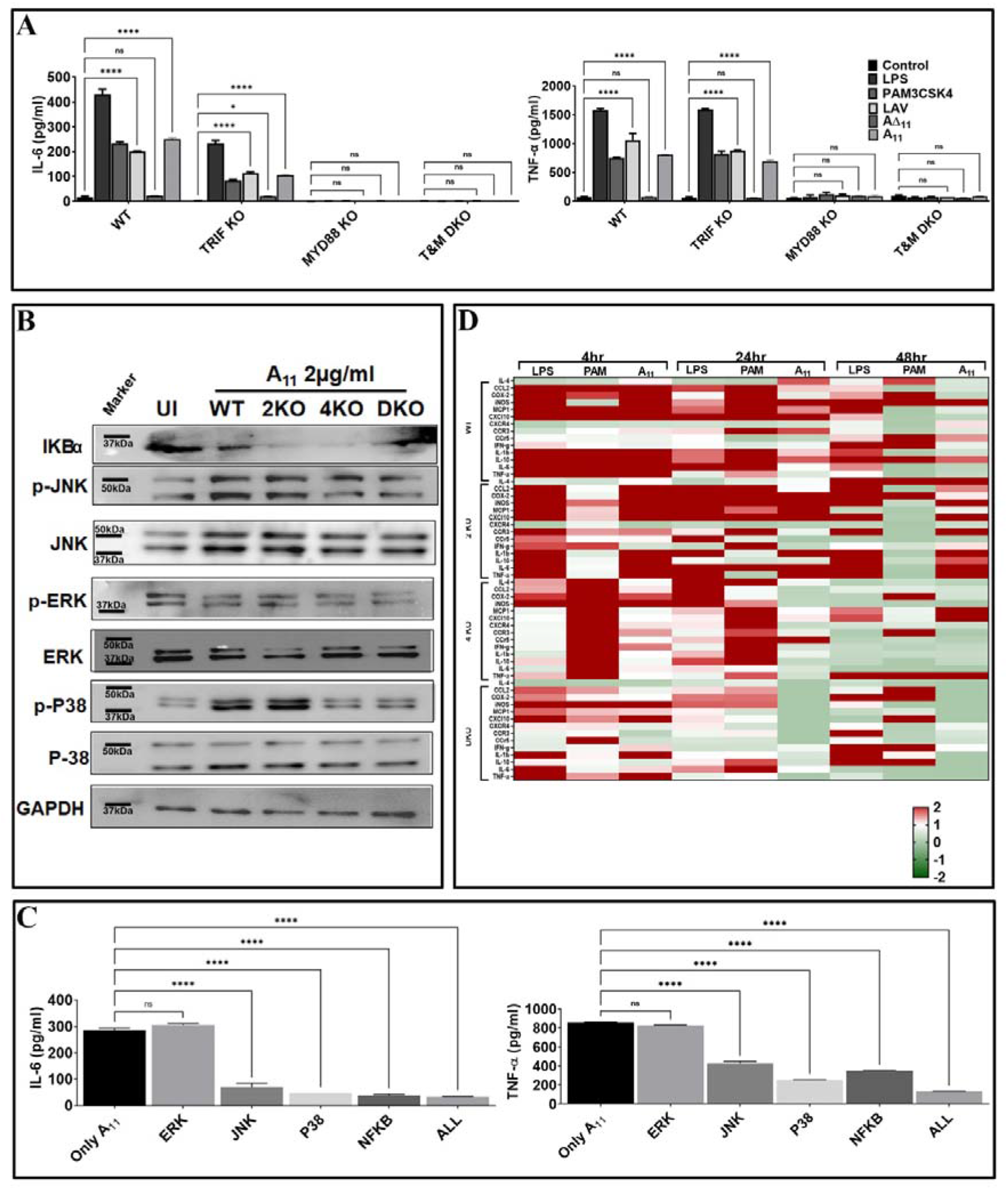
11^th^ domain of LigA (A_11_) produces pro-inflammatory cytokines via signalling through MAP kinase involving p38 and JNK pathway. **(A**) *A_11_ signals through TLR4 involving the MyD88 adapter*. WT, MyD88 KO, TRIF KO and TMDKO bone marrow-derived macrophages cell lines were treated with LAV or A_11_ or AΔ_11_ (2μg/ml) for 24h at 37°C in the presence of 5%CO_2_ and levels of IL-6 and TNF-α in the supernatants were measured by ELISA. **(B)** *A_11_ signals through TLR4 via the MAP kinase pathway involving p38 and NFkb*. WT or TLR2KO or TLR4KO or DKO macrophages cell lines were stimulated with A_11_ (2μg/ml) for 24h at 37°C in the presence of 5%CO_2_. Levels of phosphorylated p38, JNK, and ERK1/2 induced by A_11_ were analyzed by western blot as described in materials and methods. **(C)** *Pharmacological inhibitors of p38 and NFkb significantly reduces A_11_ mediated cytokine response*. RAW 264.7 cells were pre-treated for 30min with NF-kB inhibitor (SN50; 20μM), JNK inhibitor (SP600125; 40μM) or p38MAPK inhibitor (SB203580; 30μM) or ERK (U0126; 50μM) or all four inhibitors together and then stimulated with A_11_ (2μg/ml) for 24h at 37°C in the presence of 5%CO_2_ and supernatant was collected to measure levels of IL-6 and TNF-α by ELISA. **(D)** *Analysis of expression of immune response-related genes in mouse macrophages stimulated with A_11_*. WT, TLR2KO, TLR4KO and DKO mouse macrophage cell lines were treated with 2μg/ml A_11_ for 4h or 24h or 48h. Cells were recovered, RNA was isolated, converted to cDNA and gene expression was analyzed by RT-PCR as described in material and methods. The data were presented as fold changes between stimulated cells vs control and normalized to GAPDH. *E. coli* LPS (500ng/ml) or PAM3CSK4(20ng/ml) as TLR4 and TLR2 ligands respectively were used as positive controls in all experiments wherever indicated. All data are representative of three independent experiments. Significant differences were calculated using the one or two way ANOVA(****, ***, **, * indicates P < 0.0001, < 0.001, P < 0.01 and P□<□0.05 respectively).

### A_11_ is an immuno-dominant domain that induces a strong adaptive immune response

To test whether the innate response induced by A_11_ correlates to subsequent induction of adaptive response, we evaluated antigen-specific humoral, and cell-mediated immune response in mice immunized with LAV and A_11_ in alum adjuvant. Our results show that mice immunized with A_11_ with or without alum adjuvant-induced strong antibody response at day 28 post-immunization (Fig. 4A). The generation of high levels of IgG1 and significant levels of IgG2c against A_11_, indicates a mixed Th1 and Th2 response (Fig.4A). A_11_ failed to induce a significant level of IgA. A_11_ induced proliferation generation of T cells secreting significant levels of IL-4 and IL-10 (Th2 cytokines) and high levels of IFN-γ (Th1 cytokine) (Fig. 4B, 4C). There was no significant enhancement in antibody levels against A_11_ with the addition of alum adjuvant; however, cells obtained from Alum-A_11_ immunized animals produced higher levels of IL-10 and IFN-γ (Fig 4A, 4B, 4C). To test whether A_11_ is an immuno-dominant domain, we analysed IgG response at day 28 in animals immunized with LAV or A_11_ or AΔ_11_ in Freund’s adjuvant. Our results show that AΔ_11_ immunized animals induced significantly lower levels of IgG than those immunized with LAVor A_11_ (Fig.4D). This result correlated to a significant decrease in levels of IgG, cell proliferation, and induction of cytokines when serum and lymphocytes isolated from LAV-Alum immunized animals were used to analyze the response against AΔ_11_(Fig. 4E, 4F, 4G). Altogether, these results indicate that A_11_ is an immuno-dominant domain, and its deletion significantly impairs the ability of LAV to induce a robust adaptive immune response.

**Fig.4:**
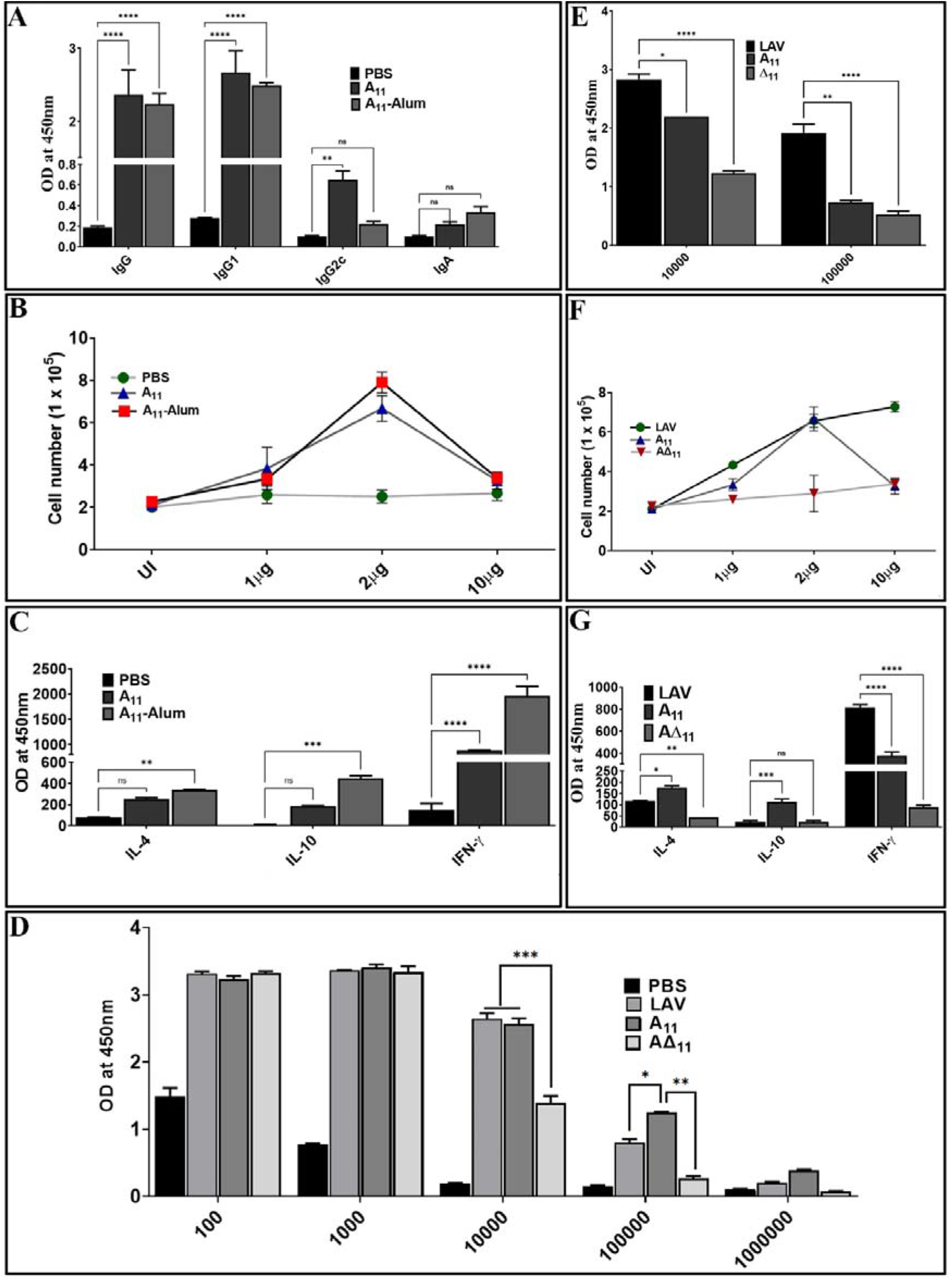
A_11_ is an immuno-dominant domain that induces robust antibody and T cell response in mice. (A) *Antibody response*. The antibody response (Total IgG, IgG1, IgG2c and IgA) on day 28 in various immunized groups (PBS, A_11_, A_11_-Alum) was evaluated by ELISA as described in materials and methods. (B) *Lymphocyte proliferation*. The proliferation of splenocytes isolated from various groups was analyzed by stimulating with recall antigen (A_11_) and counting cells after 72h. **(C)** *Cytokine analysis*. Culture supernatant from spleenocytes stimulated with A_11_ for 72h were analyzed for IL-4, IL-10 and IFN-γ by using a sandwich ELISA kit following manufacturer’s instructions. **(D)** *Total IgG response at day 28 in animals immunized with LAV, A_11_ and A*Δ*_11_ in Fruend’s adjuvant*. Total IgG at day 28 was analysed in serum from animals immunized LAV, A_11_and AΔ_11_ in Fruend’s adjuvant by ELISA as described above. **(E)** *Antibody response against A_11_ and A*Δ*_11_ in serum of animals immunized with LAV-Alum*. Total IgG was analyzed in serum from LAV immunized animals (diluted at 1:10000 and 1:100000) against A_11_ and AΔ_11_ by ELISA as described above **(F)** *Cell proliferation* and **(G)** *Cytokine analysis of lymphocytes isolated from LAV-Alum immunized animals after in-vitro stimulation with A_11_ and A*Δ*_11_*. Lymphocytes isolated at day 28 from animals immunized with LAV-Alum were stimulated with A_11_ and AΔ_11_ for 48-72h and analysed for proliferation and cytokines in the culture supernatant were determined as described above. All data are representative of three independent experiments. Significant differences were calculated using the one or two way ANOVA(****, ***, **, * indicates P < 0.0001, < 0.001, P < 0.01 and P□<□0.05 respectively).

### *Leptospira* evades complement-mediated killing by acquiring complement regulators through A_11_

*Leptospira* evades complement-mediated killing by acquiring complement regulators (FH and C4BP) or host proteases (PLG) which involves binding with surface proteins. Lig proteins, including LigA have been shown to bind to FH, C4BP and PLG. However, except for C4BP the domain/s of LigA involved in binding to FH or PLG have not been characterized. To identify and characterize the domain of LAV involved in binding to FH and PLG, we screened the individual domains (A_8_-A_13_) and corresponding deletion mutants (AΔ_8_-AΔ_13_) for their ability to bind with FH and PLG. Our dot blot result shows that only A_11_ and all the deletion mutants except AΔ_11_ were able to bind to FH and PLG (Fig. 5A). This binding was further confirmed by pulldown assay and ELISA, and our result shows that while A_11_ led to strong binding, AΔ_11_ failed to bind to both FH and PLG (Fig. 5B, 5C). We further determined if the binding of A_11_ with FH is sufficient for its functional activity. Our result shows that both LAV and A_11_ were able to bind to FH to cleave C3b in the presence of Factor I (FI) as evidenced by cleavage fragments. In contrast, AΔ_11_ failed to do so, indicating that the 11^th^ domain is involved in binding with and mediating the cofactor activity (Fig. 5D). Our ELISA result shows that A_11_ binds with PLG and converts it into active plasmin, whereas AΔ_11_ failed to generate a significant level of plasmin (Fig.5E). Western blot analysis further confirms that the released plasmin was able to cleavae C3b in presence of A_11_ whereas failed to do so in presence of AΔ^11^ indicating that 11^th^ domain is involved in PLG binding and mediating subsequent plasmin activity (Fig.5E). To establish the role of A_11_ in the complement-mediated killing, we incubated *E. coli* with 10% Normal Human Serum (NHS) pre-incubated with A_11_ or LAV or AΔ_11_. Our results show that both LAV and A_11_ domains could rescue bacteria from complement-mediated killing, but AΔ_11_ failed to do so, indicating that the 11^th^ domain is involved in evasion from complement-mediated killing (Fig. 5F, Sup. Fig.3).

**Fig.5:**
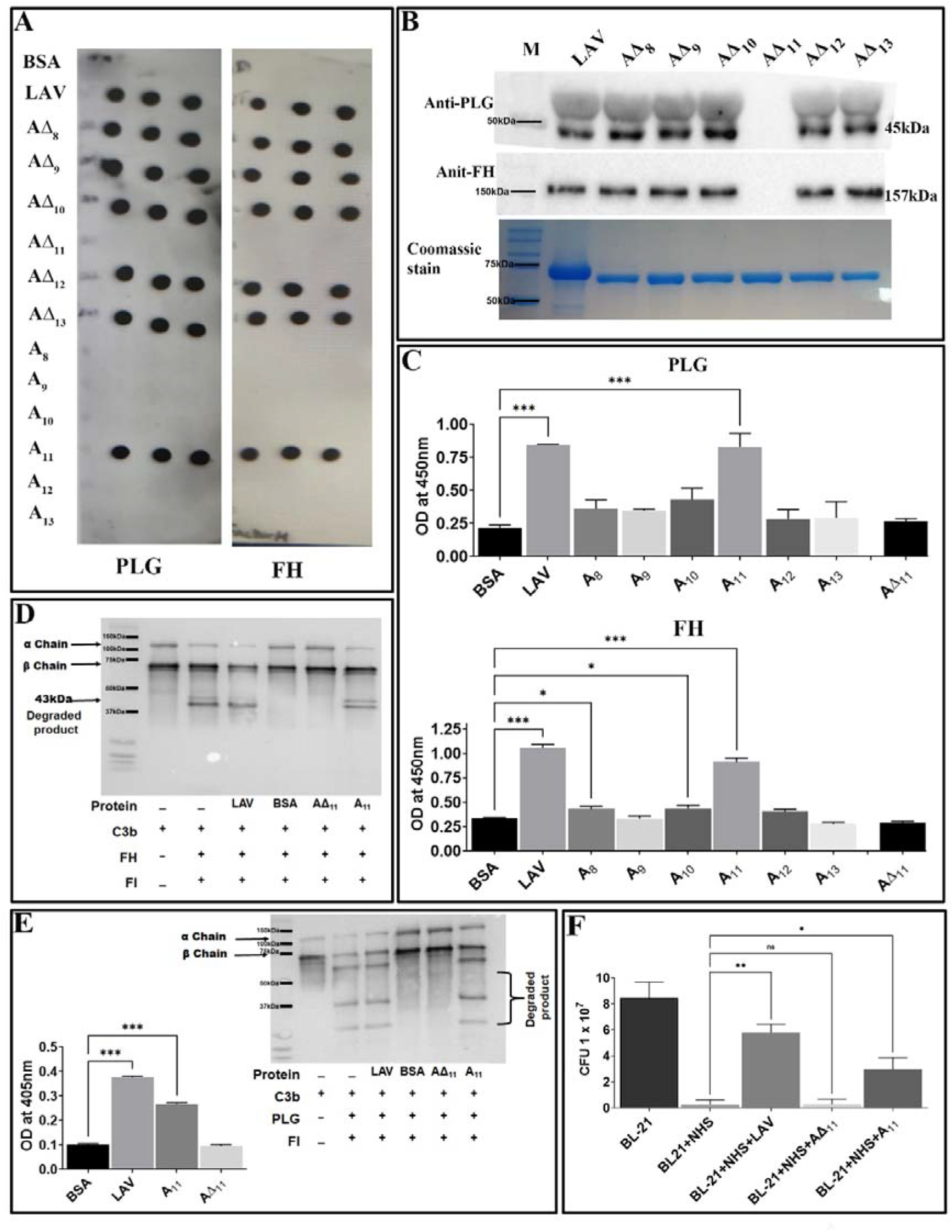
Characterization of the domain of LAV involved in evasion from complement-mediated killing. **(A)** *Binding of LAV domains as analyzed by Dot blot*. Purified proteins LAV (positive control), BSA (negative control), individual domains (A_8_-A_13_), and corresponding deletion mutants (AΔ_8_-AΔ_13_) were immobilized on nitrocellulose membranes and then incubated with 1% NHS (as a source of FH and PLG). FH and PLG were detected with specific antibodies by Western blot. **(D)** *Binding of LAV deletion mutants as analysed by Pull down assay*. Bead bound LAV deletion mutants(AΔ_8_-AΔ_13_) were incubated with 10% HI NHS and protein – protein interaction was detected by western blot using Anti FH (157 kDa) or PLG antibody (45 kDa) as described in methodology **(C)** *Binding of LAV domains as analyzed by ELISA*. Microtiter plates were coated with 1μg of proteins LAV, domains (A_8_-A_13_), and corresponding deletion mutants (AΔ_8_-AΔ_13_) and 10% HI-NHS was added to each well. The binding was detected with specific antibodies against FH and PLG as described in materials and methods. **(D)** *Co-factor activity*. LAV, A_11_, AΔ_11_ (2μg/ml), and BSA (2μg/well) were immobilized on microtiter plates and incubated with purified FH. After washing, C3b and factor I (FI) were added, and the plate was incubated for 4h at 37°C. The products were analysed by SDS-PAGE, and the cleavage fragments of C3b was detected by Western blot using anti-human C3 polyclonal antibodies as described in materials and methods. **(E)** *Plasmin activity*. LAV, A_11_, AΔ_11_ (2μg/ml), and BSA (2μg/well) were immobilized on microtiter plates followed by the addition of PLG, uPA, and specific plasmin substrate. The plate was incubated for 48h, and absorbance was read at 405nm as described in materials and methods. In another experiment, C3b was incubated with activated plasmin in the presence or absence of A_11_ and cleavage products were visualized using Western blot. **(F)** *Bactericidal assay*. 1.3□×□10^8^ *E Coli BL-2l(DE3)* cells were incubated with 10□% NHS with or without pre-incubation with A_11_ or AΔ_11_ or LAV at 20μg/ml for 30 □min at 37□°C. The samples were plated on LB agar plates and incubated at 37°C overnight. Survival was determined by counting bacterial colonies the following day. All data are representative of three independent experiments. Significant differences were calculated using the one or two way ANOVA(****, ***, **, * indicates P < 0.0001, < 0.001, P < 0.01 and P□<□0.05 respectively).

### LAV is a nuclease involved in the evasion of *Leptospira* from neutrophil extracellular traps

Recently it has been shown that *Leptospira* induces NET; however, a protein with nuclease activity to degrade NET has not been reported. To test whether LAV has nuclease activity, which might have a role in evasion from NETs, we incubated plasmid or linear DNA with varying concentrations of the protein (1 to 10μg), and our result shows that LAV was able to degrade both plasmid and linear DNA in a dose-dependent manner indicating that its having both endo and exonuclease activity (Fig 6A, 6B). To analyze if this activity is restricted to any domain, we incubated the linear DNA with individual domains (A_8_-A_13_) or deletion mutants (AΔ_8_-AΔ_13_) and our result shows that A_11_ and A_13_ were able to degrade DNA whereas all the deletion mutants except AΔ_11_ degraded DNA with equal propensity (Fig. 6C, 6D). Further, AΔ_11_ was also not able to cause significant degradation of plasmid DNA (Fig. 6E). These results indicate that LAV’s nuclease activity primarily resides in 11^th^ domain, and LAV is mediating this activity by utilizing this domain. To test whether LAV can cleave the Neutrophil Extracellular Trap (NET) we stimulated the mouse neutrophils with PMA to induce NET and then treated them with LAV (5μg/ml). Our confocal microscopy result shows that LAV could degrade the PMA induced NET, further confirming its nuclease activity and possible role in degrading NETs *in vivo* (Fig.6F). These results indicate that LAV has nuclease activity that *Leptospira* might exploit to evade from NETosis.

**Fig.6:**
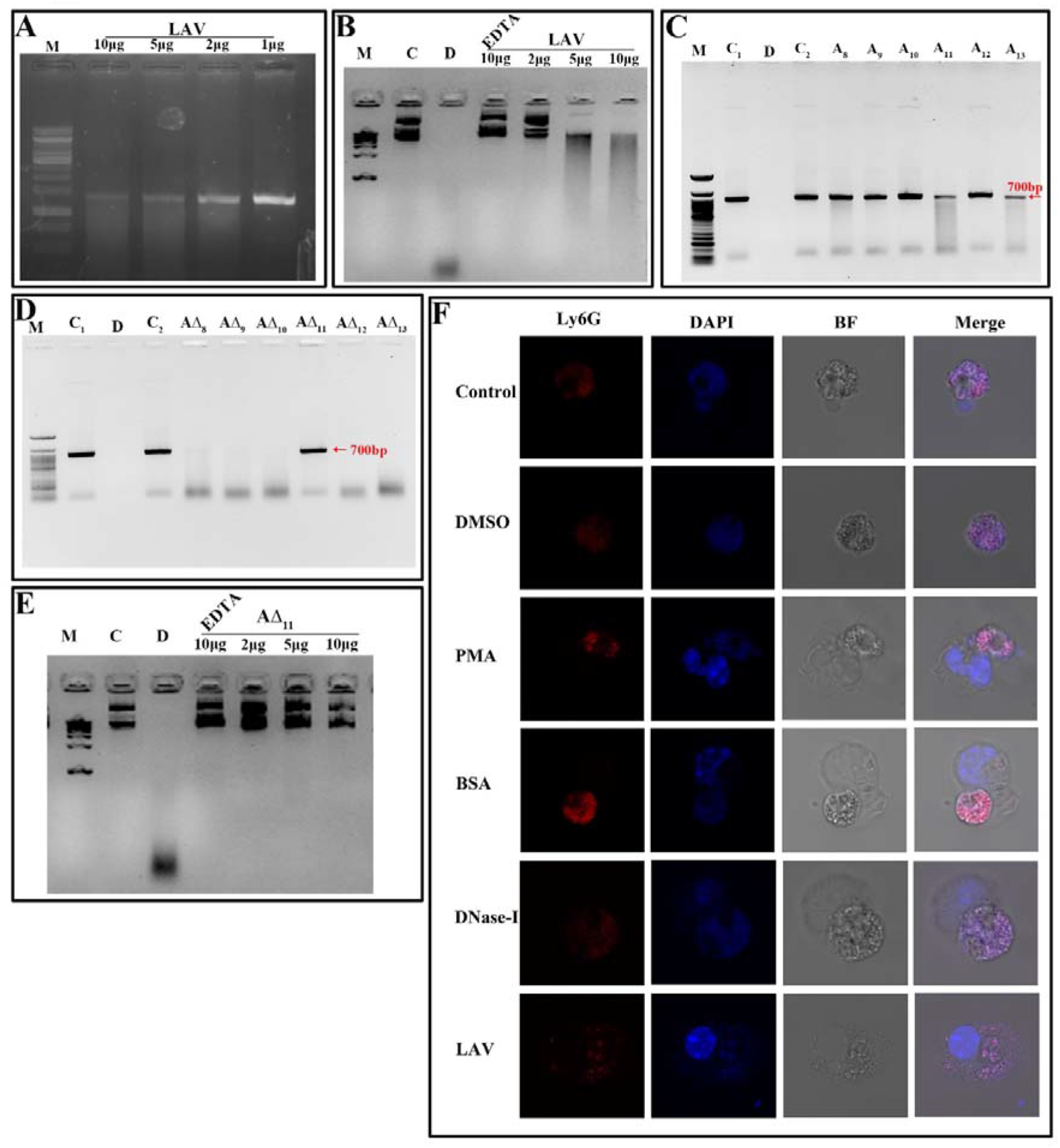
LAV is nuclease capable of degrading Neutrophil Extracellular Trap (NET) **(A)** *Exonuclease activity of the LAV*. 700bp DNA (200ng) was incubated with different concentrations of LAV (2, 5 and 10μg) in DPBS with 5mM MgCl2 at 37°C for 3h followed by visualized using the Agarose gel electrophoresis.**(B)** *Endonuclease activity of the LAV*. Plasmid DNA (200ng) was incubated with different concentrations of LAV (2, 5 and 10μg) in DPBS with 5mM MgCl2 at 37°C for 3h followed by visualized using the Agarose gel electrophoresis. **(C)** *Exonuclease activity of the LAV domains*. 700bp DNA (200ng) with incubated with various LAV domains (A_8_-A_13_) at 5μg in DPBS with 5mM MgCl2 incubated at 37°C for 3h followed by EtBr-Agarose gel electrophoresis. **(D)** *Exonuclease activity of domain deletion mutants of LAV*. DNA (200ng) with incubated with various domain deletion mutants of LAV (AΔ_8_-AΔ_13_) at 5μg in DPBS with 5mM MgCl_2_ incubated at 37°C for 3h followed by EtBr-Agarose gel electrophoresis **(E)** *Endonuclease activity of the* AΔ_11_. Plasmid DNA (200ng) was incubated with different concentrations of AΔ_11_ (2, 5 and 10μg) in DPBS with 5mM MgCl_2_ with or without EDTA at 37°C for 3h followed by visualized using the Agarose gel electrophoresis. C and C1 indicates DNA alone, D indicatres DNA treated with DNAse, C2 indicates DNA with reaction mixture, M is 100bp or 1kb DNA ladder in all experiments wherever indicated **(F).** *NETosis assay:* Mouse Neutrophils were cultured on glass coverslips stimulated with DMSO or LPS (500ng) or PMA (0.5μM) for 3.5h and then treated with DNase-I (positive control) or BSA (negative control) or LAV (şig) for 2h at 37°C and visualized under 63X of Leica microscopy. DAPI; staining of the complete DNA content (Nuclear and released), Ly6G; neutrophil marker, BF; Bright field. All data are representative of three independent experiments.

## Discussion

The ability of *Leptospira* to cause persistent infection and efficient colonization in a variety of hosts reflects its potential to subvert or thwart the innate immune response^15^. This ability has been attributed to the procession of a wide variety of surface molecules like proteins, lipopolysaccharide, etc., which are redundant in their function and may also undergo structural variation to avoid recognition by the host immune system been observed in other spirochetes ^12^. Surface proteins, including lipoproteins from spirochetes like *Borrelia* and *Treponema* play a critical role in immune evasion by limiting their expression or inducing antigenic variation after infection, which greatly enhances host infectivity and persistence^13,14,45^. *Leptospira*, like other pathogens, may voluntarily interact with TLRs (or other innate receptors) through surface molecules (proteins, LPS) but might evade this recognition through multiple mechanisms to establish infection or fitness in the host^46–48^. Both TLR2 and TLR4 receptors play a major role in host defense against *Leptospira* infection^21^. It has been shown that TLR4 plays a critical role in controlling bacterial load and developing severe leptospirosis in mice ^49^. Thus, it is likely that those surface proteins and LPS which are natural ligand of these receptors and can activate macrophages and DCs, might modulate their expression or undergo variations thereby enabling the bacteria to evade this innate recognition as has been reported for other pathogens. Several surface proteins of *Leptospira* have been identified as ligands of TLR2 or TLR4 capable of activating the innate immune response and are potential vaccine candidates ^22,23,50–52^. Lig proteins are important virulence factors, and their expression during infection or loss of expression during *in vitro* culture has been correlated to virulence of the infecting serovar ^34,53^. These proteins interact with various host molecules, including extracellular matrix (ECM), coagulation cascade, and complement regulators. LigA is the most promising vaccine candidate. It has been demonstrated that the variable region of the protein comprising domains 10-13 (LigA10-13) is sufficient in inducing protection against challenge in the hamster model^36–38,40^. Thus, the diverse functions of LigA prompted us to investigate whether their role is limited to binding to host ECM and complement regulators, or they are also involved in the modulation of the host innate immune response, thereby contributing to infection and persistence in the host.

Since several investigators have established the protective role of a variable region of LigA (LAV), we chose to decipher its role in the modulation of the host innate immune response. We cloned, expressed and purified the recombinant LAV and tested its ability to activate mouse macrophages. Our result shows that LAV could activate macrophages, as evident from the production of pro-inflammatory cytokines (Fig.1). This effect was not due to contaminating LPS as pretreatment with Polymixin B didn’t attenuate, whereas digestion with Proteinase K abrogated the cytokine production (sup Fig.1B). The LAV-induced activation was TLR4 dependent as evident from its binding to the receptor, induction of IL-8 by TLR4 transfected HEK293 cells, and abrogation of cytokine production in TL4KO mouse macrophages. We expected that LAV might not signal through TLR2 as it is devoid of the signal sequence and hence lipidation. However, several non-acylated proteins like LcrV from *Yersinia*, MPB83, and PPE18 from *Mycobacterium*, PorB from *Neisseria* and FimA from *P. gingivalis* have been shown to signal through TLR2 ^54–58^. Although rare, several proteins from other bacterial pathogens have been reported to induce TLR4 dependent production of pro-inflammatory cytokines and expression of surface markers ^59,60^. Further, lipidated recombinant proteins, which usually signal through TLR2 due to the lipid moiety, may signal through TLR4 if unlipidated as has been observed in the case of Omp16 and Omp19 of *Brucella* ^61,62^. Moreover, recombinant unlipidated rBCSP31 from *Brucella abortus* and rLsa21 from *Leptospira* have been shown to signal through both TLR2 and TLR4 and induce activation of macrophages^22,63^. Since LAV is composed of several immunoglobulin-like repeat domains, we attempted to identify the domain involved in innate immune activation. Our results shows that A_11_ is involved in TLR4 dependent activation of mouse macrophages leading to production of pro-inflammatory cytokines and expression of costimulatory molecules and maturation marker (Fig. 2 and 3). Further, A_11_ modulated the expression of several innate responses related to genes (cytokines, chemokines, and surface receptors) involved in the activation and maturation of macrophages (Fig.3). Our results are in accordance with previous reports, where several TLR4 ligands, including bacterial proteins, have been shown to activate macrophages and DCs via signalling through the MAP kinase pathway, leading to the induction of cytokines, expression of surface markers, and immune response-related genes^59,64–67^. Since TLR, dependent activation of innate response, is essential for T cell expansion, differentiation, and memory formation, we tested the adaptive response induced by A_11_ in mice. Our result shows that the strong innate response induced by A_11_ also correlated to the generaion of higher level of adaptive response (Fig. 4). Interestingly the response generated against A_11_ was equivalent to LAV and was significantly decreased in terms of antibody titer, cell proliferation, and cytokines in the absence of this domain, indicating that A_11_ is the most immuno-dominant domain capable of inducing robust antibody and T cell response. Although A_11_ induced mixed Th1 and Th2 response, high levels IFN-γ may be correlated to strong activation of innate response, particularly innate B cells, which require investigation^68^. The strong adaptive response induced by A_11_ without any adjuvant highlights its immunomodulatory potential. It suggests that TLR4 dependent signaling by A_11_ might activate strong innate and subsequent adaptive response leading to clearance of *Leptospira* from the host. We speculate that to evade this protective response, *Leptospira* might limit expression or undergo antigenic variation in LigA to avoid recognition with TLR4 and subsequent activation of the innate and adaptive response. However, this needs to be tested, and experiments are ongoing. Further, the inability of AΔ_11_ to activate a strong innate and subsequent adaptive response is not due to misfolding, as mutant protein retains significant numbers of alpha-helix and beta sheets as revealed by structural analysis (sup Fig. 2).

It is known that pathogenic *Leptospira* is resistant to the bactericidal activity of normal human serum (NHS)^69,70^. They can evade complement attack by using various strategies like recruitment of the host complement regulators, acquisition of host proteases or secretion of proteases that can cleave complement components on the bacterial surface and in its surroundings^8^. Several surface proteins of *Leptospira* like LenA, LenB, LcpA, Lsa30, including Lig proteins (LigA and LigB) have been shown to bind to various complement regulators ^20,25–30,42,43,71–74^. Moreover, both conserved and the variable (N and C terminal) regions of LigA and LigB are involved in binding to FH and C4BP. Our results confirmed the previous report of binding of LAV with FH and PLG and identified and characterized the domain/s involved in binding and mediating subsequent co-factor or plasmin activity (Fig. 5). Additionally, the rescue of *E. coli* from complement-mediated killing in NHS pre-incubated with A_11_ further substantiates the critical role of this domain in complement evasion^75^ (Fig.5F). Thus, binding of FH and PLG to A_11_ reflects the ability of *Leptospira* to utilize this domain for simultaneously inhibiting lectin and alternate pathway of complement-mediated killing (Fig. 5 and sup Fig.4). To our knowledge, this is the first report that demonstrates that a single domain of a surface protein is alone capable of recruiting FH and PLG directly from NHS and prevents complement activation.

Apart from killing bacterial pathogens by intracellular ROS and phagocytosis, neutrophils might release neutrophils extracellular traps (NETs) that capture and kill microbes in the extracellular space in tissues (at sites of infection) or within blood vessels^17,76^. This mechanism on killing extracellular bacteria by trapping outside the cell is independent of phagocytosis and degranulation ^17^. Several bacterial pathogens, including *Staphylococcus aureus, Clostridium perfringens* and *Streptococcus pyogenes* have evolved sophisticated mechanisms to suppress, escape, and/or resist NETs through surface proteins having nuclease activity^77–79^. Recently, it has been shown that *Leptospira* can induce the NET, and its surface protein LipL21 can modulate neutrophil function; however, nuclease capable of degrading NET has not been reported^16,31^. Our study discovered the nuclease (DNase) activity of LAV and demonstrated that A_11_ primarily or predominantly mediates this activity as it was able to degrade DNA with the same propensity as LAV (Fig. 6). Our results further demonstrate that LAV or A_11_ exhibits both endo and exonuclease activity. Although nuclease activity of LAV is not restricted to A_11_, as significant activity was also mediated by A_13_ but our result clearly shows that it is primarily mediated by A_11_. The ability of LAV to degrade PMA induced NET in mouse neutrophils and highlights the possible role of LigA in escaping the bacteria from NETosis. To our knowledge, this is the first report of identification of nuclease activity of a surface protein in *Leptospira* and also demonstrating its diverse role in modulating the host innate response.

In conclusion, our results demonstrate that LigA is a multifunctional protein involved in attachment to host cells to initiate infection, a TLR4 agonist which can activate a strong innate response (possibly evading this TLR4 activation by antigen variation or downregulating its expression upon infection in the host), binds to complement regulators to evade complement-mediated killing and exhibit nuclease activity when *Leptospira* gets entrapped in NET (Fig. 7). These features might contribute to its successful colonization in a particular host. Interestingly, these functions are mediated primarily by a single domain (A_11_) which lies in LAV. This promising vaccine candidate conferred protective immunity against lethal infection in the hamster model of the disease. Thus, the protective efficacy of LAV based vaccine may be correlated with its ability to induce the robust antigen-specific humoral and T cell response that might lead to the generation of antibodies conceivably blocking binding to host extracellular matrix, acquiring complement regulators and inhibiting DNase activity all of these may aid in the clearance of bacteria from the host (Fig. 7). Our results provide important insight into the role of LAV in host-pathogen interaction and also established it as an immuno-modulator or adjuvant, which makes it an ideal candidate for developing vaccines for this dreadful zoonosis^80^.

**Fig.7.**
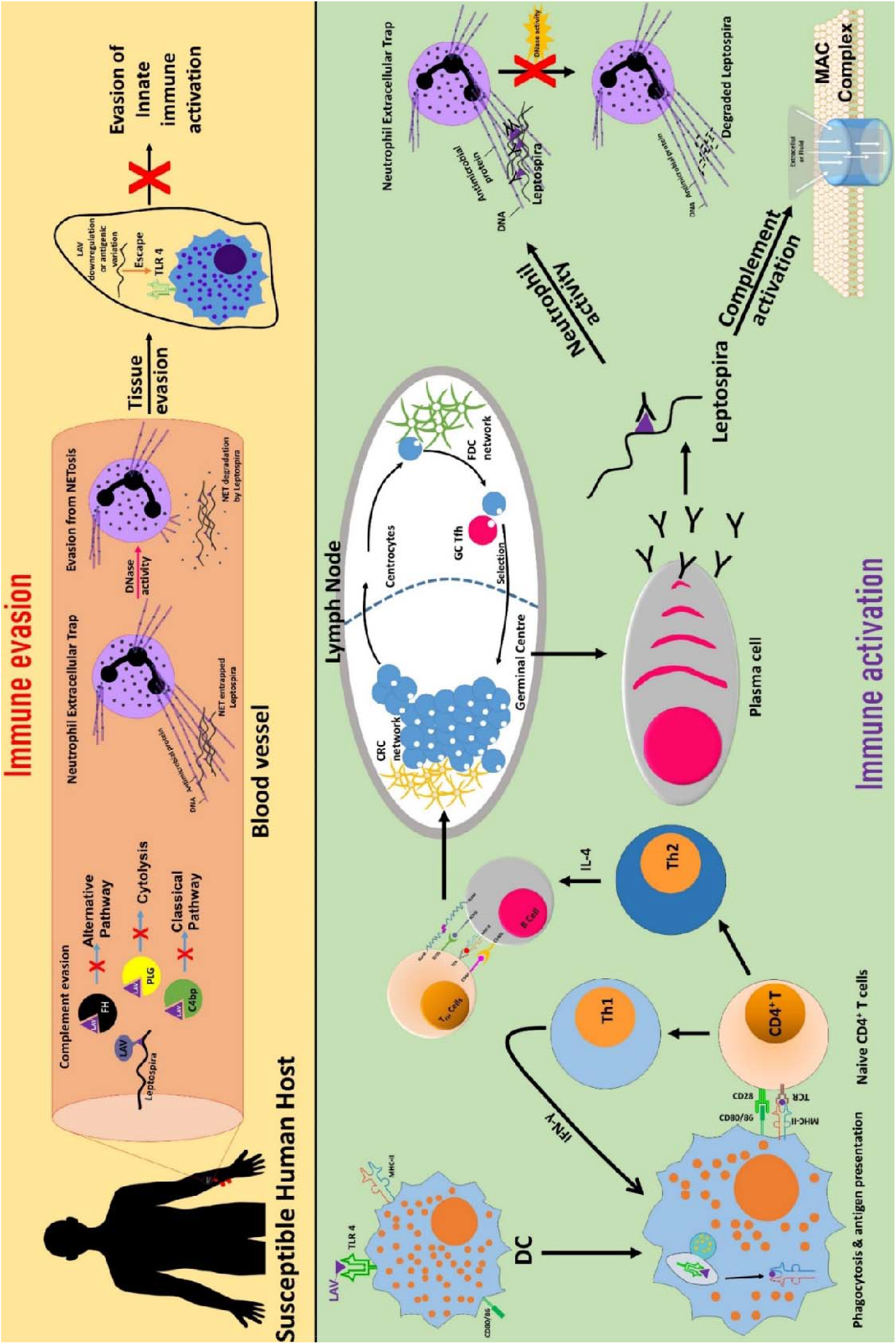
Schematic presentation of the role of LigA in the modulation of host immune response. **(A)** *Immune evasion*. LigA expressed during infection might acquire complement regulators (FH, C4BP, PLG) to inhibit both the classical and alternate pathways of complement-mediated killing. *Leptospira* might utilize the nuclease activity of LigA to escape from NET. Upon interaction with host innate immune cells (DCs, macrophages), LigA might undergo antigenic variation or downregulate its expression to evade recognition through TLR4 and subsequent activation of the innate response. **(B)** *Immune activation*. LigA (LAV) can activate strong innate and adaptive immune responses leading to the production of antibodies that may block binding to complement regulators, inhibit nuclease activity and enhance phagocytosis, all of which may contribute to the killing of bacteria and clearance from the host.

## Material & Methods

### Animals, cell lines and reagents

Male C57BL/6J mice (6-8 weeks) were obtained from the Animal Resource and Experimental Facility of NIAB, Hyderabad. The original breeding colonies were obtained from Jackson Labs, USA. The animals were maintained in a pathogen-free condition. All the procedures for animal experiments were approved by the Institutional Animal Ethics Committee (IAEC) and performed in accordance with the Committee for the Purpose of Control and Supervision of Experiments on Animals (CPCSEA) guidelines. RAW264.7 and HEK293 cell lines were originally purchased from the ATCC (Manassas, VA). Mouse macrophage WT (NR-9456), TLR2KO (TLR2-/-, NR-9457), TLR4KO (TLR4-/-, NR-9458), DKO (TLR2-/-/4-/-, NR-19975), TRIFKO (TRIF-/-, NR-9566), MyD88KO (MyD88-/-, NR-15633) and TMDKO (TRIF-/-MyD88-/-, NR-15632), cell lines were obtained from BEI Resources, USA. Cells were cultured in DMEM (Sigma, USA) supplemented with 10% FBS (Invitrogen, Carlsbad, CA, USA), penicillin (100 U/ml), and streptomycin (100 μg/ml) and maintained at 37°C in a humidified incubator (5% CO2). Pharmalogical inhibitors of NF-kB (SN50), p38 (SB203580), ERK (U0126) and JNK (SP600125) were purchased from Invivogen. Mouse IL-6 and TNF-α Sandwich ELISA kits were from R&D Biosystems. APC onjugated hamster antimouse-CD80, PE-conjugated rat antimouse-CD86, BV421 conjugated rat antimouse-CD40 and Per Cp Cy5.5 conjugated rat antimouse-MHC-II antibodies were procured from BD biosciences, US. Normal Human Serum (S1-100ml), Goat anti FH (SAB2500260), Mouse anti PLG (SAB1406263-50UG), Plasmin substrate, UPA (SRP6273) complement C3b (204860-250G) Complement factor I (C5938-1MG), Complement factor H (C5813-1MG), plasminogen (SRP6518-1MG) were procured from Sigma Chemical Co, USA. Polyclonal anti C3 was purchased from Complement Technology, USA.

### Cloning, expression, and purification of recombinant proteins

The Lig A variable (LAV) gene sequence was amplified by PCR from *L. interrogans* serovar Pomona strain genomic DNA using specific primers and then cloned in His-Sumo tagged pET28a expression vector. Domains of LAV 8 to 13 and corresponding deletion mutants (AΔ_8_-AΔ_13_) were similarly cloned in the pET28a vector. Various domain deletion mutants of LAV (AΔ_8_-AΔ_13_) were generated by PCR-based site-directed mutagenesis. All the clones were verified by sequencing. The plasmid was transformed into *BL21(DE3)* Rosetta. The resulting transformants were grown at 37°C overnight on LB broth containing 50μg/ml kanamycin, and the expression of the protein was induced with 1 mM isopropyl β-D-1-thiogalactoside (IPTG). The cells were harvested by centrifugation at 10,000 rpm, and the cell pellet was resuspended in 100mM Tris HCl, 150mM NaCl pH8.0, followed by sonication at constant pulses. The lysate was centrifuged to remove cell debris, and the supernatant was subjected to affinity chromatography using Ni-NTA beads column (Takara). Eluted protein was dialyzed against 1×PBS with four changes for two days at 4°C. The protein was then passed through Detox-Gel (Pierce, USA) to remove any contaminating LPS from *E. coli*, and a residual trace amount of LPS was monitored by Limulus amoebocyte lysate (LAL, Endotoxin Detection Kit, Pierce, Thermo, USA) assay following the manufacturer’s instructions. The purified protein was checked for size and purity by SDS-PAGE, and concentration was estimated using the Bradford reagent (Sigma, USA).

### Cell stimulation assays by Cytokine ELISAs

Cytokine ELISA kits (R&D systems) were used to measure cytokine levels, following the manufacturer’s instructions. RAW264.7 cells were stimulated with LAV or (A_8_-A_13_) or corresponding deletion mutants (ΔA_8_-AΔ_13_) (2μg/ml), PAM3CSK4 (20ng/ml), and *E. coli* LPS 0111-B4 (500ng/ml) for 24h at 37°C in the presence of 5%CO_2_ and cytokines (IL-6, TNF-α) were measured in the culture supernatant according to the manufacturer instructions. The proteins were pre-treated with Polymyxin B (20ng/mg protein) at 37°C for 1hr and proteinase K (5μg/mg protein) at 65°C for 1hr followed by inactivation at 95°C for 5min before each assay to rule out endotoxin activity. In a separate experiment wild type, TLR2KO, TLR4KO, DKO, MyD88KO, TRIFKO, TMDKO macrophage cell lines were stimulated with PAM3CSK4, LPS, LAV or A_11_ or AΔ_11_ for 24h at 37°C/5%CO_2_ and cytokines (IL-6, TNF-α) in the culture supernatant were measured by ELISA kit as per manufacturer’s instructions. HEK-293T cells were cultured in a complete DMEM medium for 24h at 37°C in the presence of 5%CO_2_ and transfected with TLR2, TLR4, and NF-kB reporter plasmids using X-fect Transfection reagent (Takara, Japan) following manufacturer’s protocol. Cells were stimulated with LAV (2μg/ml) for 24h, and then IL-8 levels were measured in the cell culture supernatant. To assess the signaling pathway involved, additional experiments were done in which RAW264.7 cells were pre-treated for 30min at 37°C/5%CO_2_ with pharmacological inhibitors of NF-kB (SN50; 20μM) or JNK (SP600125; 40μM) or p38MAPK (SB203580; 30μM) or ERK (U0126; 20μM) followed by treatment with A_11_ (2μg/ml) for 24h at 37°C in the presence of 5%CO_2_. Cytokine levels were measured by ELISA kit.

### Flow cytometric analysis

RAW264.7 cells were incubated in 6-well plates (0.3×10^6^ cells/well) with PAM3CSK4 (20ng/ml), LPS (500ng/ml), LAV or A_11_ or AΔ_11_ (2μg/ml) for 24h at 37°C in the presence of 5%CO_2_. Cells were harvested and washed with pre-chilled PBS and then incubated on ice for 1h in the dark with respective fluorochrome conjugated antibodies against CD80, CD86, MHC-II, and CD40. Cells were washed and then fixed with 1% paraformaldehyde, and 50,000 total events/sample were acquired using a BD Fortessa. The data were analyzed using FlowJo software.

### Preparation of antisera

Male C57BL/6J mice (6-8 weeks) were immunized subcutaneously on days 0 with 20μg of LAV, A_11_, AΔ_11_ in complete Freund’s adjuvant (CFA) and then boosted on day 21 with 10μg of proteins in Incomplete Freund’s adjuvant (IFA). Sera were collected one week after booster (day 28) and titer were determined using ELISA. The mouse serum having anti-LAV, anti-A_11_ or anti-AΔ_11_ antibodies were used in confocal microscopy.

### TLR binding assay

WT, TLR2KO, TLR4KO and DKO cell lines were grown overnight on glass-bottom cell imaging dishes (Eppendorf) and then incubated for 30 min at 37°C in the presence of 5%CO_2_ with LAV or A_11_ or AΔ_11_ (2μg/ml) in DMEM without FBS. The cells were washed with PBS and fixed for 15 min using 4% paraformaldehyde followed by blocking with 5%FBS in PBS for 30min at RT. The cells were then incubated with anti-LAV or A_11_ or AΔ_11_ (mouse serum, 1:100 dilution) for 1h, washed three times with PBS, and then stained with Alexa Flour 647 conjugated rabbit anti-mouse IgG (Biolegend, USA). Cells were extensively washed and mounted with VECTA SHIELD (containing DAPI) mounting medium and observed using a 63x oil objective on a confocal microscope (Leica SP8, Wetzlar, Germany).

### RT-PCR

WT, TLR2KO, TLR4KO and DKO mouse macrophage cell lines were treated with A_11_ (2μg/ml), LPS (500ng/ml) or PAM3CSK4 (20ng/ml). After 4, 24 and 48h of treatment cells were recovered in 500μl of TRIzol (Invitrogen, Carlsbad, CA), and equal volumes of chloroform were added; samples were centrifuged at 12000□rpm for 15□min at 4°C. The aqueous phase was then passed through RNA easy mini columns (MN) and RNA was purified following the manufacturer’s protocol. RNA quality was checked by running on a Formaldehyde gel for 18s and 28s RNA bands and analyzed on Bioanalyser. The RNA quantity was assessed by UV spectroscopy and purity by 260/280 ratio. First-strand cDNA was synthesized using the superscript III-RT system (Invitrogen) following the manufacturer’s instructions. RT-PCR was performed in 96 well microtiter plates in a 10μl reaction volume containing 50ng cDNA, 10μM each primer (Table 1) and SYBR green (Bio-Rad). Samples were run in triplicate, and data was analyzed with Sequence Detection System (Bio-Rad CFX-96). The experimental data were presented as fold changes of gene expression of stimulated cells at various time points relative to control. mRNA levels of the analyzed genes were normalized to the amount of GAPDH present in each sample.

**Table 1.**
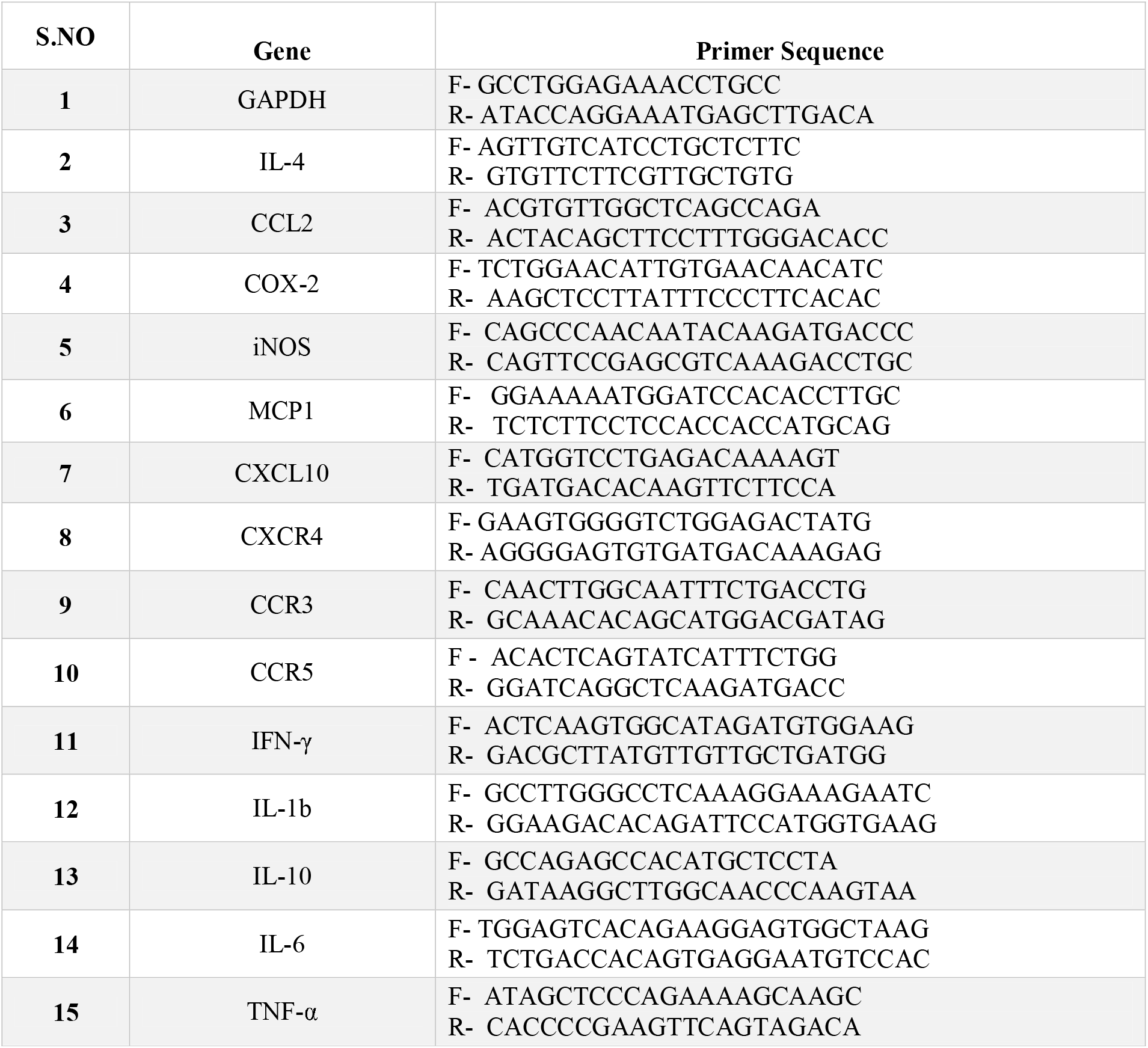
Primers used for RT-PCR.

**Table 2.**
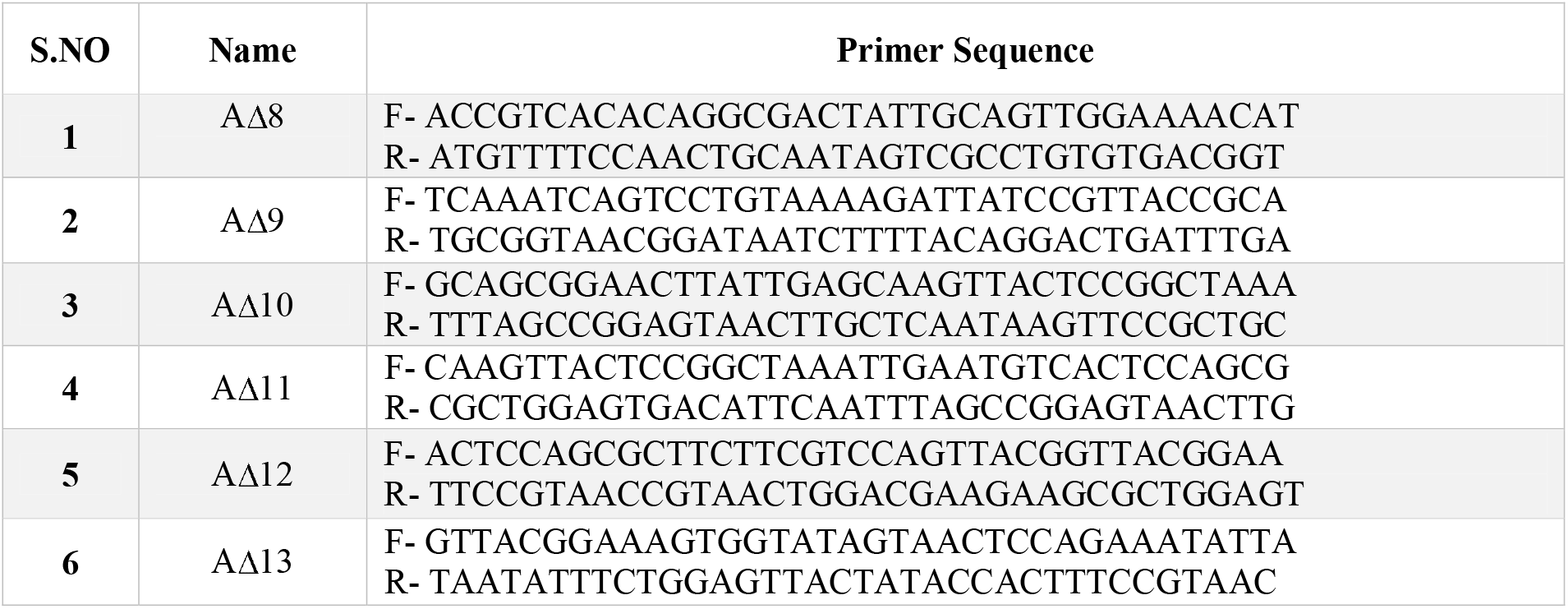
Primers used for domain deletions.

### Circular Dichroism

The proteins (LAV and AΔ_11_) were dialysed against sodium phosphate buffer and the CD spectroscopy of the far-UV spectrum was obtained in a Jasco J-810 spectropolarimeter (Japan Spectroscopic). The resulting spectra are presented as the averages of three scans recorded from 190 to 260 nm. The residual molar ellipticity is expressed in degree cm2 dmol-1. Spectral data were analyzed with the software BESTSEL (https://bestsel.elte.hu/) for estimation of the secondary structure content.

### Adaptive immune response

Male C57BL/6J mice (6-8 weeks) were immunized subcutaneously on days 0 with 20μg and on day 21 with 10μg of LAV or A_11_ with or without Alum adjuvant. Animals vaccinated with PBS were used as control. Mice bled at various time points a (day 0, 21 and 28), and the serum was analyzed for antigen-specific antibodies. Animals were euthanized on day 28 and blood, and spleens were collected for evaluation of antigen-specific immune responses. To determine antibody response, serum samples from individual mice were collected on the day before immunization and then on day 21 and 28. Total IgG, IgG1, IgG2a, and IgA concentrations were evaluated using ELISA using the standard procedure. For cell proliferation assay, splenocytes prepared from different mice groups were stimulated with varying concentrations (1, 2 and 10 μg/ml) of LAV or A_11_ or AΔ_11_. Cells were counted under an inverted microscope at 24h, 48h and 72h post-stimulation. To determine Th1 and Th2 cytokines, culture supernatant was collected at 48-72h post antigen stimulation and were used to estimate the IL-4, IL-10 and IFN-γ using cytokine ELISA kits (R&D systems) following the manufacturer’s instructions.

### Dot Blot binding Assay

Dot blot binding assays were performed to confirm the binding of various LAV domains (A_8_-A_13_) and their single domain deletion mutants (AΔ_8_-AΔ_13_) with FH and serine protease PLG. 1μg of each protein (wild type, single domain, and domain deletion mutant) was immobilized onto NC membranes (0.2μ pore size; Bio-Rad). The membranes were kept for drying for 5-10 min at RT. The membranes were then blocked with 5% BSA in Tris-buffered Saline-Tween 20 (TBS-T) for 2h at RT, washed three times with TBS-T, and incubated with 1% normal human serum (NHS) diluted in PBS with gentle shaking for 3h at RT. After extensive washing with TBS-T, the membranes were incubated with the corresponding primary antibody (Goat anti-FH, Mouse anti-PLG 1: 10,000 dilution) in TBS-T for 2h at RT. The membranes were then washed with TBS-T and incubated with a respective peroxidase-conjugated secondary antibody (1: 6,000) for 2h at room temperature. Reactive spots were developed using a chemiluminescence system with an exposure time of 10 sec.

### Pull down assay

Each domain deletion mutants of LAV (AΔ_8_-AΔ_13_) along with 10% heat-inactivated NHS, were incubated with 15 μL of Ni-NTA agarose beads (Takara) overnight at 4°C. Agarose beads were washed five times with PBS containing 40mM imidazole and then interacting proteins were eluted with PBS containing 250 mM imidazole. Each elutes boiled in reducing Laemmli buffer and subjected to SDS-PAGE. The proteins were transferred to nitrocellulose membrane followed by western blot against Anti-FH, and Anti-PLG, respectively.

### ELISA binding assay

Protein binding to soluble complement regulators FH, C4BP and PLG were analysed by ELISA as described previously ^27,29^. Briefly, micro titre plates were coated overnight at 4°C with 1μg of domains (A_8_-A_13_) and their single domain deletion mutants (AΔ_8_-AΔ_13_). BSA and LAV were used as positive and negative control. The wells were washed three times with PBS containing 0.05 % Tween 20 (PBS-T), blocked with 300μl PBS/3 % BSA for 2h at 37°C, and incubated with 100μl 10 %NHS for 90 min at 37°C. After washing with PBST, goat anti-FH (1:1000), rabbit anti-C4BP (1:1000) or mouse anti-PLG (1:5000) was added and the plate was incubated for 1h at 37°C. After washing, HRP-conjugated anti-goat or anti-mouse or anti-rabbit IgG was added and incubated for 1h at 37°C. The wells were washed and TMB substrate was added (100μl/well). The reaction was stopped by the addition of 50μl 2N H2SO4 and absorbance was measured at 450nm in a microplate reader.

### Cofactor activity assay

Cofactor activity was determined as described previously^29^. Briefly, 2μg of A_11_, AΔ_11_ and LAV (positive control) or BSA (negative control) were coated on microplates overnight at 4°C. The wells were washed and blocked with PBS/2 % BSA for 2h at 37°C, followed by the addition of 2μg pure FH and further incubation for 90 min at 37°C. Unbounded FH was removed by washing and then 250ng FI and 500ng C3b were added to the microtiter plate wells and incubated for 3-5h at 37°C. The supernatants were loaded onto a 10% SDS-PAGE gel and transferred onto a 0.22μ PVDF membrane. For the immunoblotting, membranes were blocked with 5 % BSA and then incubated with goat anti-human C3 (1:5000) for 2h at RT. After the usual steps of washing, the membranes were incubated with a peroxidase-conjugated secondary antibody. The images were visualized under the Clarity Max Western ECL substrate (BIO-RAD) using Syngene G: BOX Chemi XX6/XX9.

### Plasmin activity assay

Plasmin activity was determined as described previously^29^. Briefly, Microtiter plate wells were coated overnight at 4°C with 2μg LAV (positive control), BSA (negative control), A_11_, AΔ_11_. The plate was washed with PBS-T and blocked for 2h at 37°C with 10% skim milk. After discarding the blocking solution, human PLG (2μg/well) was added, followed by incubation for 90 min at 37°C. After washing plates three times with PBS-T, 250μg/ml uPA was added with the plasmin-specific substrate, *D*-Val-Leu-Lys 4-*p*-nitroanilide dihydrochloride (100μl/well) at a final concentration of 0.4mM in PBS. Plates were incubated for 24h at 37°C, and absorbances were measured at 405nm using a microplate reader.

### Bactericidal assay

Bactericidal activity was determined as described elsewhere^29^.1.3□××10^8^ *E Coli (BL21DE3)* cells were washed once with PBS and incubated with 10□% NHS with or without pre-incubation with recombinant proteins (A_11_, AΔ_11_, LAV at 20μg/ ml) in a final reaction volume of 100μl for 30□min at 37°C. The samples were placed on ice to stop further bacteriolysis and then plated on LB agar plates. The plates were incubated at 37°C overnight. Survival was determined by counting bacterial colonies the following day.

### Nuclease activity

To examine the DNase activity of LAV, 200ng of 700bp DNA was incubated with different concentration (1 or 2 or 5 or 10μg) of LAV or 2μg domains (A_8_-A_13_) and 2μg domain deletion mutants (AΔ_8_-AΔ_13_) or DNase I (20IU, positive control) in DPBS with 5mM MgCl2 in a PCR tube at 37°C for 2h. The reaction mixture was subjected to EtBr Agarose gel electrophoresis (1%) and then observed under the Gel doc. In a separate experiment, a plasmid was used as a substrate to check endonuclease activity of various concentrations (2ug, 5ug, and 10ug) of rLAV or rAΔ_11_. EDTA is a nuclease activity inhibitor, was used as control.

### Isolation of Neutrophils from murine bone marrow

Neutrophils were isolated from the bone marrow of C57BL/6J mice using the standard procedure^44^. Briefly, bone marrow was flushed from femurs using a 26G needle, passed through a 30μm cell strainer, and then cells were washed in complete RPMI-1640 twice at 1,400 rpm for 10 min at 4°C. After lysis of RBCs using ACK lysis buffer, cells were washed with RPMI-1640 supplemented with 10% FBS, counted, and resuspended in 1ml of ice-cold sterile PBS. Next, cells were overlaid on 3ml of Histopaque 1077/1119 mix in a 15ml conical tube and then centrifuged for 30min at 2,000 rpm at 25°C without braking. Neutrophils at the interface were collected and washed twice with a complete RPMI-1640 medium, counted and suspended in the medium for the specific assay. The viability was determined by Trypan blue exclusion assay.

### NET assay

2 x 10^5^ freshly isolated neutrophils in 300μl medium were added to the imaging dish and kept at 37°C in the presence of 5%CO_2_ overnight. Cells were treated with 3μl of DMSO or PMA (50ng/ml) or LPS (100ng/ml) and further incubated for 3h at 37°C/5%CO_2_. Cells were washed thrice with DPBS and then incubated with LAV (2μg/ml) or DNase-I (20IU) or BSA (5μg/ml) in 5mM MgCl2 containing PBS for 2h at 37°C/5%CO_2_. Cells were washed with DPBS, fixed with 4% PFA (15min at RT), and then stained with Rat anti-mouse Ly6G (Alexa flour 647) for 30min. Cells were washed thoroughly with DPBS, mounted with VECTA SHIELD (with DAPI) mounting medium and observed using a 63x oil objective on a confocal microscope (Leica SP8, Wetzlar, Germany).

### Statistical Analysis

For all the experiments, wherever required one way or two-way ANOVA were executed to analyze the results. The data were represented as the mean of triplicates□±□SEM. p□<□0.05 was considered significant.

## Data Availability

The data that support the findings of this study are available from the corresponding author upon reasonable request.

## Acknowledgments

This work was supported partly from the DBT project-BT/PR21430/ADV/90/246/2016 (SP025) on developing *Leptospira* vaccines and partly from DBT-NIAB flagship project No-BT/AAQ/01/NIAB-Flagship/2019 (SP051) on host-pathogen interaction, which are funded to SMF from the Department of Biotechnology, Ministry of Science and Technology, Government of India. Financial support from the NIAB core fund is duly acknowledged. The authors would like to thank the Director, NIAB, Dr. Subeer S. Majumdar, for providing the necessary infrastructural facility and support for the execution of the above study. Thanks to Mr. Shashikant Gawai and Mrs. Rama Devi for helping in confocal microscopy and Flow cytometry. Thanks to Dr. Jayant Hole for help in animal experiments. AK is supported by UGC fellowship and registered for Ph.D. program at RCB, Faridabad. VPV is supported by CSIR fellowship and registered for Ph.D. program at Manipal University, Manipal.

## Author’s contribution

SMF conceived the idea and designed the experiments. AK, VPV, SK, MA, PV, MAT performed the experiments. AK, VPV, SK and SMF analyzed the data. VPV made all the figures of the manuscript and did the statistical analysis. YFC contributed reagents and input in experiment design and data analysis. SMF, AK, and VPV wrote the initial draft, and SMF edited the manuscript. All authors approved the final version of the manuscript.

## Supplementary Materials

**Sup. Fig. 1:**
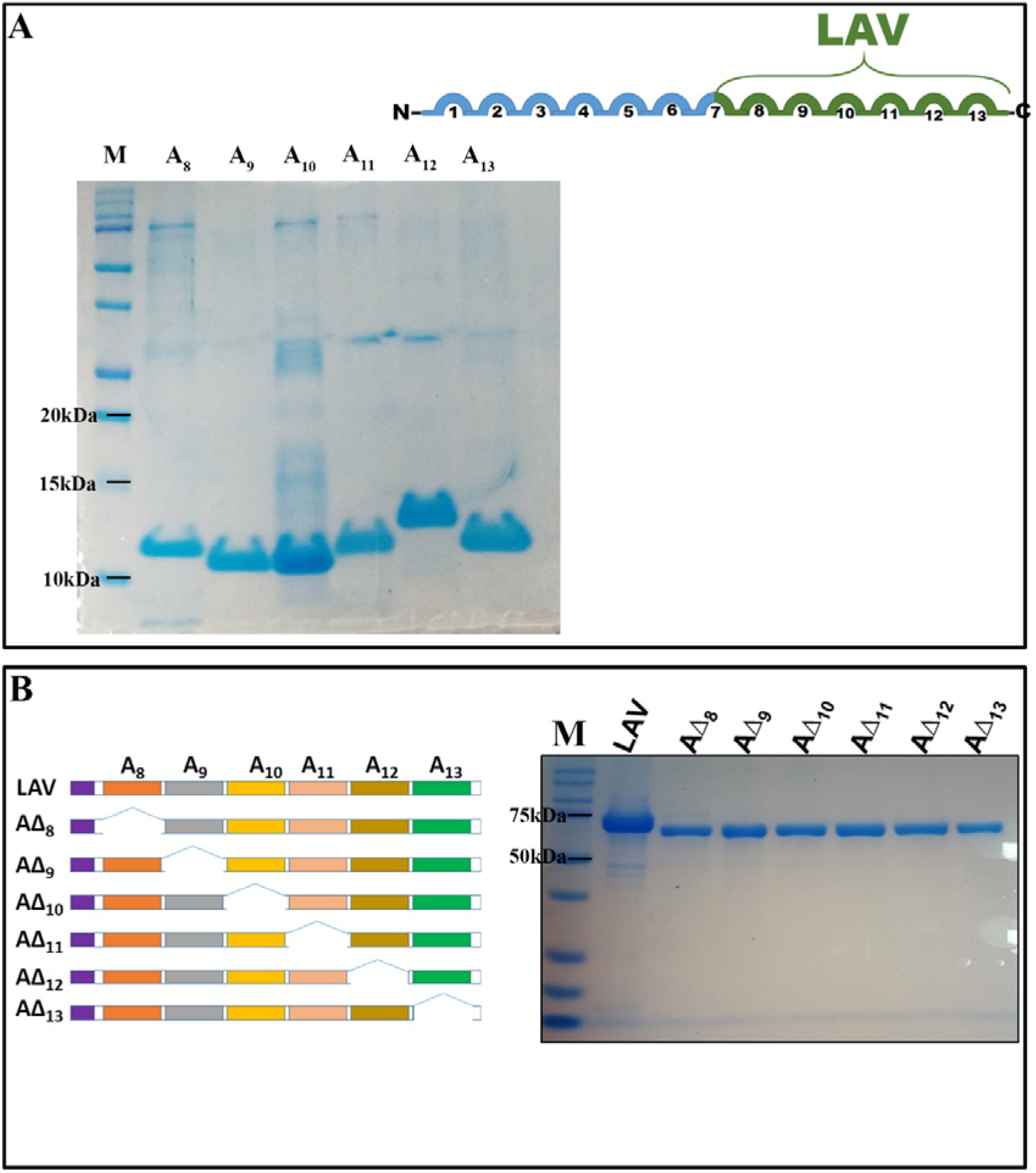
Purification of recombinant proteins. **(A)** *SDS PAGE profile of domains of Variable A (LAV)*. The recombinant proteins were purified as His-sumo fusion proteins as described in materials and methods. The expected molecular weight of each domain ranges from 11-12kd. **(B)** *Schematic presentation of strategy of creating LAV domain deletion mutants (AΔ_8_-AΔ_13_) by PCR based site-directed mutagenesis and SDS PAGE profile of purified proteins*. The recombinant proteins were purified as His-sumo fusion proteins as described in materials and methods The expected molecular weight of LAV was 73kd and each domain deletion mutant was ~63kd. Data are representative of three independent experiments. (*Indicates P□<□0.05).

**Sup. Fig. 2:**
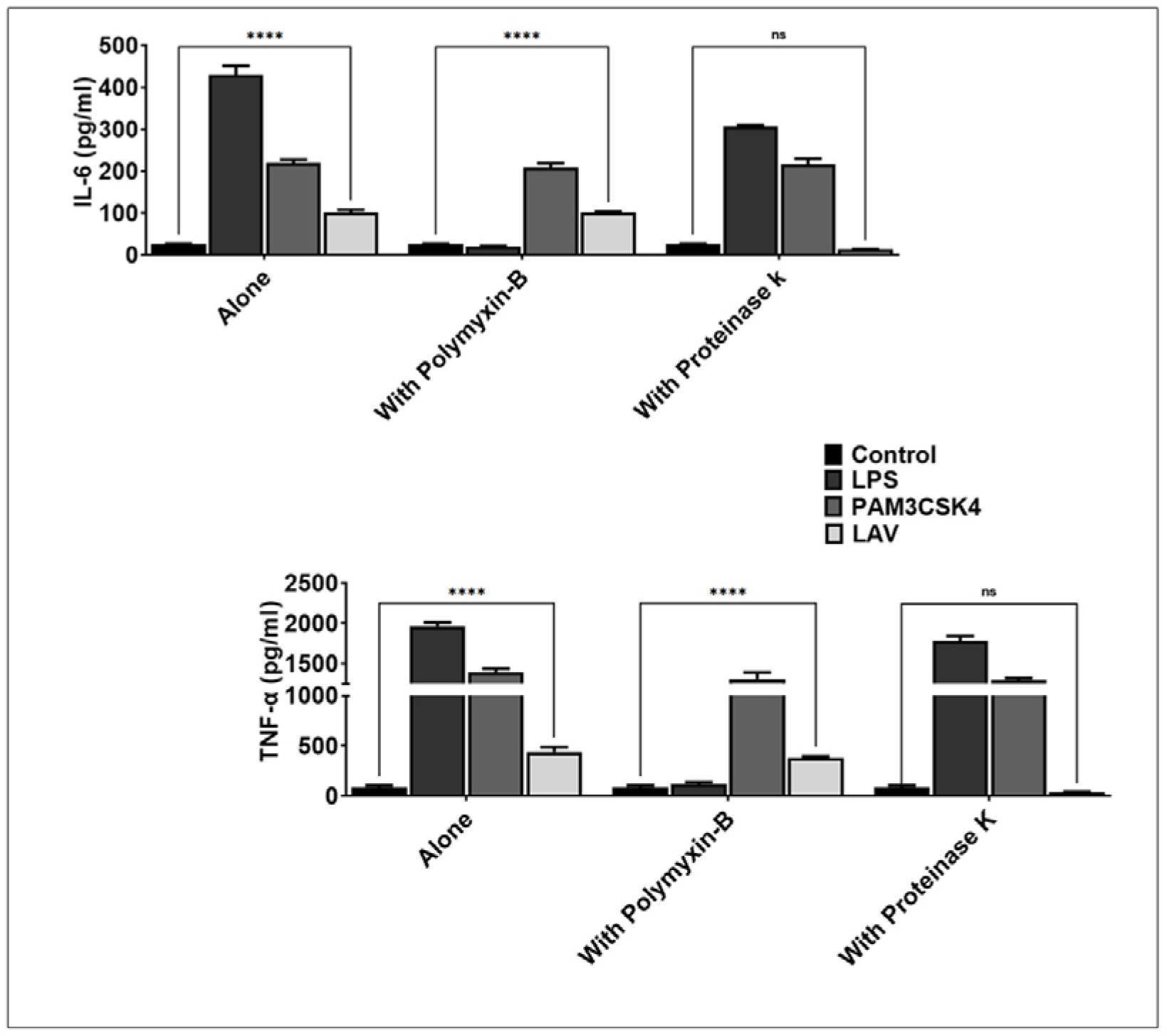
Effect on TLR activity after pre-treatment of purified recombinant proteins with Polymixin B and Proteinase K. RAW264.7 cells were incubated with 2μg/ml of purified LAV pre-treated with Polymyxin B or Proteinase-K as mentioned in materials and methods and supernatant was collected to measure levels of IL-6 and TNF-α by ELISA.

**Sup Fig. 3.**
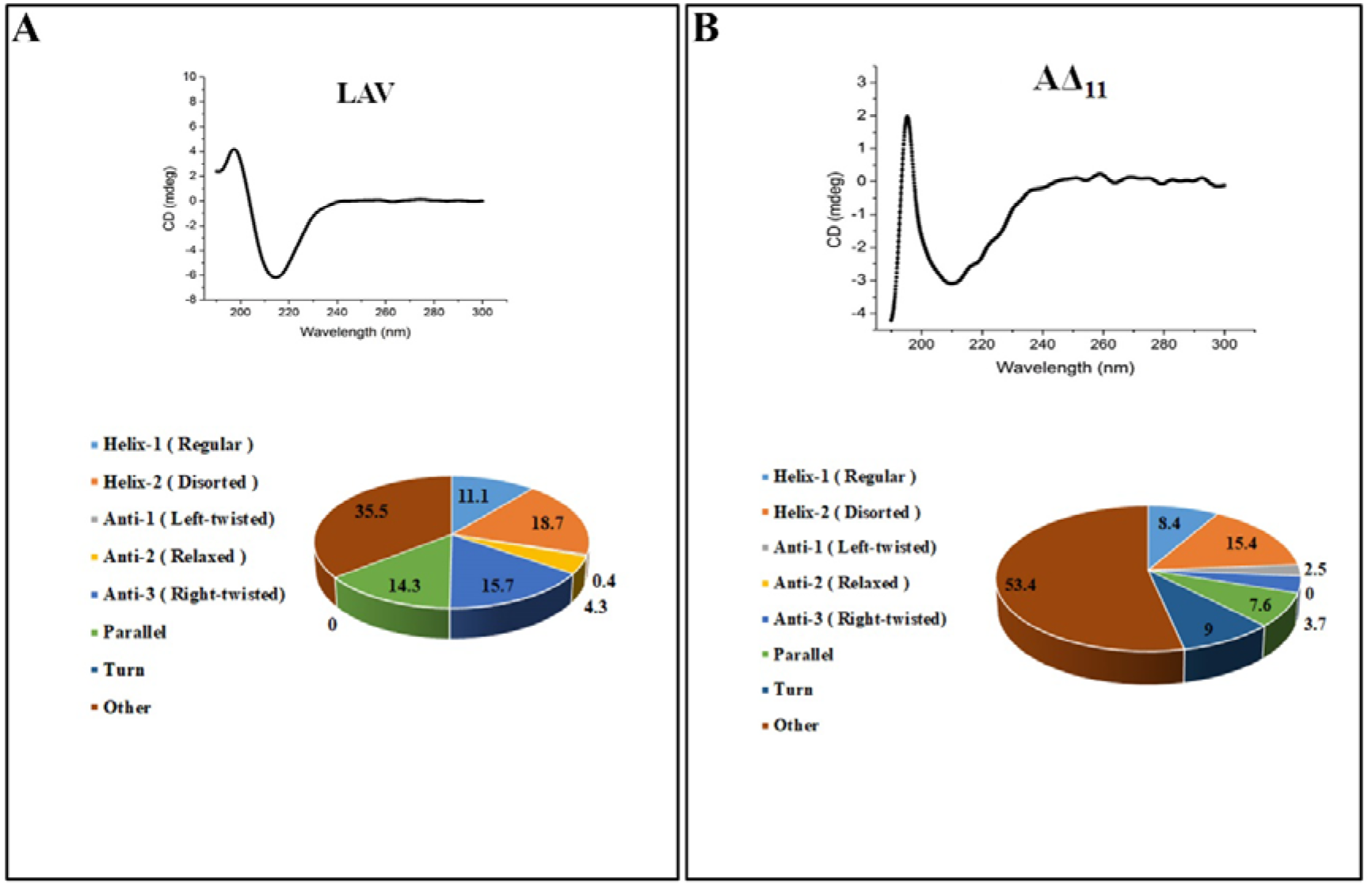
Circular dichroism spectra of the recombinant proteins. CD spectra of recombinant proteins LigA WT and LigAΔ_11_. Far-UV CD spectra are presented as an average of five scans recorded from 190 to 300 nm.

**Sup Fig. 4.**
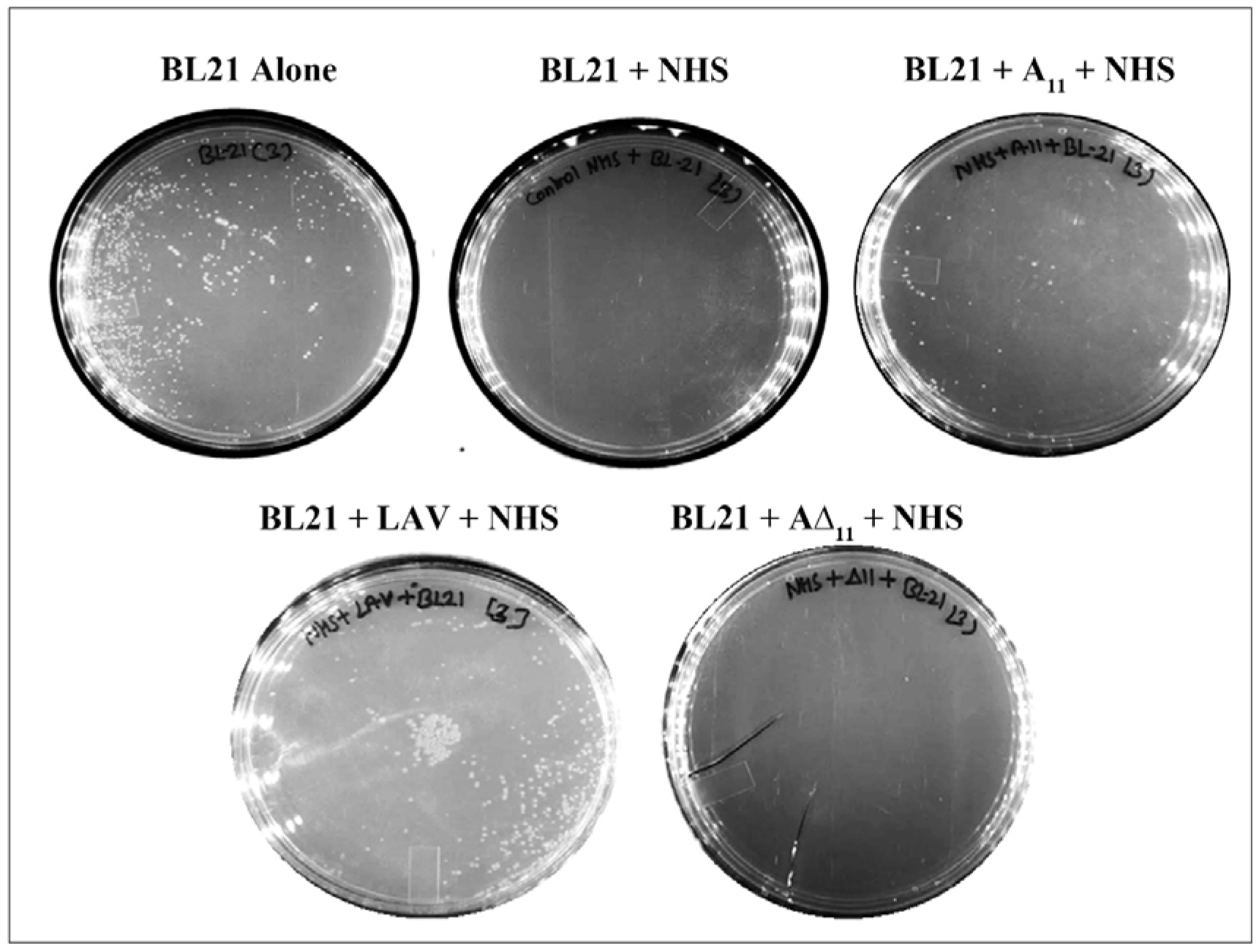
Bacterial survival assay. Photographic images of survival assay bacterial colonies formed by BL-21 *E. coli* after the treatment with NHS, which is pre-incubated with LigAvWT, Δ11 and A_11_.

